# Getting Personal with Epigenetics: Towards Machine-Learning-Assisted Precision Epigenomics

**DOI:** 10.1101/2022.02.11.479115

**Authors:** Alex Hawkins-Hooker, Giovanni Visonà, Tanmayee Narendra, Mateo Rojas-Carulla, Bernhard Schölkopf, Gabriele Schweikert

## Abstract

Epigenetic modifications are dynamic control mechanisms involved in the regulation of gene expression. Unlike the DNA sequence itself, they vary not only between individuals but also between different cell types of the same individual. Exposure to environmental factors, somatic mutations, and ageing contribute to epigenomic changes over time, which may constitute early hallmarks or causal factors of disease. Epigenetic changes are reversible and, therefore, promising therapeutic targets. However, mapping efforts to determine an individual’s cell-type-specific epigenome are constrained by experimental costs. We developed eDICE, an attention-based deep learning model, to impute epigenomic tracks. eDICE achieves improved overall performance compared to previous models on the reference Roadmap epigenomes. Furthermore, we present a proof of concept for the imputation of personalised epigenomic measurements on the ENTEx dataset, where eDICE correctly predicts individual- and cell-type-specific epigenetic patterns. This case study constitutes an important step towards robustly employing machine-learning-based approaches for personalised epigenomics.

---

Epigenetic mechanisms play an essential role in developmental biology and human disease [1, 2]. They act at the intersection of genetic and environmental factors to control, regulate, and propagate cellular responses, significantly contributing to diverse cellular phenotypes. Importantly, their influence on gene activity is reversible without altering the underlying DNA sequence. They, therefore, provide unique diagnostic and therapeutic opportunities and offer promising targets for precision medicine approaches [3–5], with particular interest for applications in cancer treatment [6, 7]. However, crucial challenges remain, mainly because epigenomes are cell-type specific and dynamically changing on different time scales - for example during the cell cycle, development, or ageing. Therefore, decoding epigenetic patterns is particularly laborious, expensive, and data-intensive.

A deeper understanding of epigenetic modifications has shed new light on the mechanisms involved in certain neurological and neurodegenerative diseases, developmental disorders, and some forms of cancer [3, 8–10]. Large-scale efforts to map the functional properties of human epigenomes proved essential for these developments and have provided a crucial resource to understand how the interplay between genetic and epigenetic factors affects cellular identity and function [11, 12]. While these projects aim to profile diverse cell types using various epigenetic assays comprehensively, the experimental costs impose constraints that inevitably lead to incomplete maps, with many cell types still sparsely analysed.

As a result, computational approaches that can leverage existing epigenomic data to impute the results of as-yet unperformed assays are of considerable interest. To this end, imputation models seek to exploit the correlations between sets of epigenomic marks within and between cell types to predict unobserved signal values across the genome. The challenge the imputation problem presents is one of combinatorial generalisation: given existing genome-wide measurements for a set of combinations of tissue or cell type and experimental assay, these methods seek to generate genome-wide predictions for the combinations of cell type and assay for which experimental measurements are currently unavailable.

## Results

### eDICE and previous work on epigenomic imputation

In 2015, Ernst and Kellis pioneered work in the field of large-scale epigenomic imputation by introducing ChromImpute [13], offering exciting possibilities for epigenomic profiling at scale, quality control, improvement of downstream analysis, and the optimisation of biological experiments [13, 14]. ChromImpute employs a regression framework that continues to show strong performance in accurately predicting unobserved epigenomic tracks. Despite its excellent performance, ChromImpute’s design presents two significant drawbacks. First, it requires the training of a new ensemble model for each target cell-assay combination, which is a severe computational constraint when considering the dimensionality of complete personalised and cell-type specific epigenomic maps [15]. Second, ChromImpute’s use of track-specific feature sets means information is not readily shared between predictive models for different tracks, limiting the extent to which it is able to exploit the increasingly large amount of data gathered by consortia such as ENCODE [16].

Overcoming these limitations, other imputation models have recently been developed which are based on factorisation frameworks and can generate genome-wide predictions for arbitrary combinations. Here, the complete set of possible epigenomic measurements (i.e. the set of all possible combinations of cell line, epigenomic assays and genomic locations) are represented as a single data tensor, and missing entries of this tensor are reconstructed via combinations of learned vector embeddings (‘factors’) representing the index along each of the dimensions. In particular, PREDICTD [17] trains an ensemble of linear tensor factorisation models, in which learned vectors representing cell line, assay type and genomic position are combined linearly via a generalised inner product to produce predicted values. Subsequently, Avocado [18] introduced the use of a neural network to output a nonlinear combination of the cell, assay and position embeddings, leading to improved performance. While the simplicity and relative parsimony of these approaches is appealing, and the ability to share information between tracks suggests greater potential to exploit the full array of available epigenomic measurements, the performance of these methods is competitive with ChromImpute on only a subset of metrics [18]. Furthermore, the requirement to independently learn embeddings to represent each position in the human genome, even at 25 base-pair resolution, imposes daunting memory demands.

We propose a novel method, eDICE (***e****pigenomic* ***D****ata* ***I****mputation through* ***C****ontextualized* ***E****mbeddings*), based on a new formulation of the epigenomic imputation problem designed to combine the strengths of the regression and tensor factorisation frameworks. Similar to regression approaches, eDICE uses the signal values of observed tracks at the target position as inputs. However, by using these signal values to build separate representations of (i) the local epigenomic state of each cell and (ii) the local enrichment of each mark, eDICE also adopts the principle of factorised representation learning which underlies the combinatorial generalisation capacity of tensor factorisation approaches. At the same time, we remove the need to learn explicit embeddings of genomic position. The rationale behind this is that genomic locations can implicitly be grouped through shared functional properties, e.g. inaccessible intergenic regions or active promoters, enhancers. These shared local properties are reflected in the partially observed marks, which we use directly to infer the remaining marks. In contrast, PREDICTD and Avocado use the observed marks to explicitly learn individual genomic embeddings along the complete genome, which are subsequently exploited for the actual imputation task.

### A single compact model with reduced memory requirements

The goal of imputation methods is to predict the signal values *y*^(*k*)^(*c, a*) for epigenomic tracks generated by performing assay *a* in cell type *c* at all binned genomic locations *k*. To achieve combinatorial generalisation we seek to learn an imputation function having the factorised form 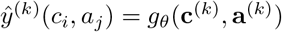, where **c**^(*k*)^ and **a**^(*k*)^ are local representations of the cell type and assay, reflecting their properties at the *k*^*th*^ genomic bin, and *g*_*θ*_ is a neural network. The crucial difference from previous tensor factorisation models is that this function operates on local rather than global representations of cell types and assays, thereby avoiding the need for explicit representations of each genomic location. Following this factorised form, there remains the problem of how to produce the relevant embeddings.

Intuitively, we would like the local cell type embeddings to reflect the latent epigenomic state of each cell type and, similarly, the assay embeddings to reflect a latent ‘profile’ summarising the signal observed when applying the assay across different cell types at the given genomic location. To infer these latent representations, we use the measured local signal values in each cell type and assay to produce embeddings that are a function not only of this cell type or assay-specific signal but also of the observed *context* of measurements in related cell types and assays. For example, a cell type in which a measurement of a particular histone modification is missing might then nonetheless be associated with a representation that encodes whether similar cell types carry that histone modification at the position in question. The problem is thus viewed as producing *contextualised embeddings* for each of a set of cell types and each of a set of assays, which is the kind of set-based representation learning problem for which self-attention and in particular the Transformer architecture has been shown to provide a powerful inductive bias [19, 20]. We therefore use separate Transformer-style networks to map the local signal associated with each cell type and each assay to sets of contextualised local cell type and assay embeddings, which are thus conditioned on the local signal in all observed tracks (i.e. the set of values *y*^(*k*)^(*c, a*) for all observed pairs (*c, a*)). To produce a prediction, the contextualised cell type and assay embeddings corresponding to the desired output track are combined via the network *g*_*θ*_ for which, following Avocado, we use a multi-layer perceptron (MLP). Full details of the architecture are provided in the Online Methods and schematised in Figure 1a. We note that as a result of not requiring learned embeddings for all genomic bins, our model’s parameter count is significantly smaller than other single-model imputation approaches based on full tensor factorisation (Table 1).

**Table 1.** Number of models and parameters per model required to make genome-wide predictions on the Roadmap test set. While PREDICTD and Avocado require several billion parameters for genome-wide prediction, eDICE requires only 6 million to obtain competitive performance.

| Model | # models | # parameters per model |
| --- | --- | --- |
| ChromImpute | 203 | / |
| PREDICTD | 8 | $\sim 11.5 B$ |
| Avocado | 1 | $\sim 3.4 B$ |
| eDICE | 1 | $\sim 6 M$ |

**Fig. 1.**
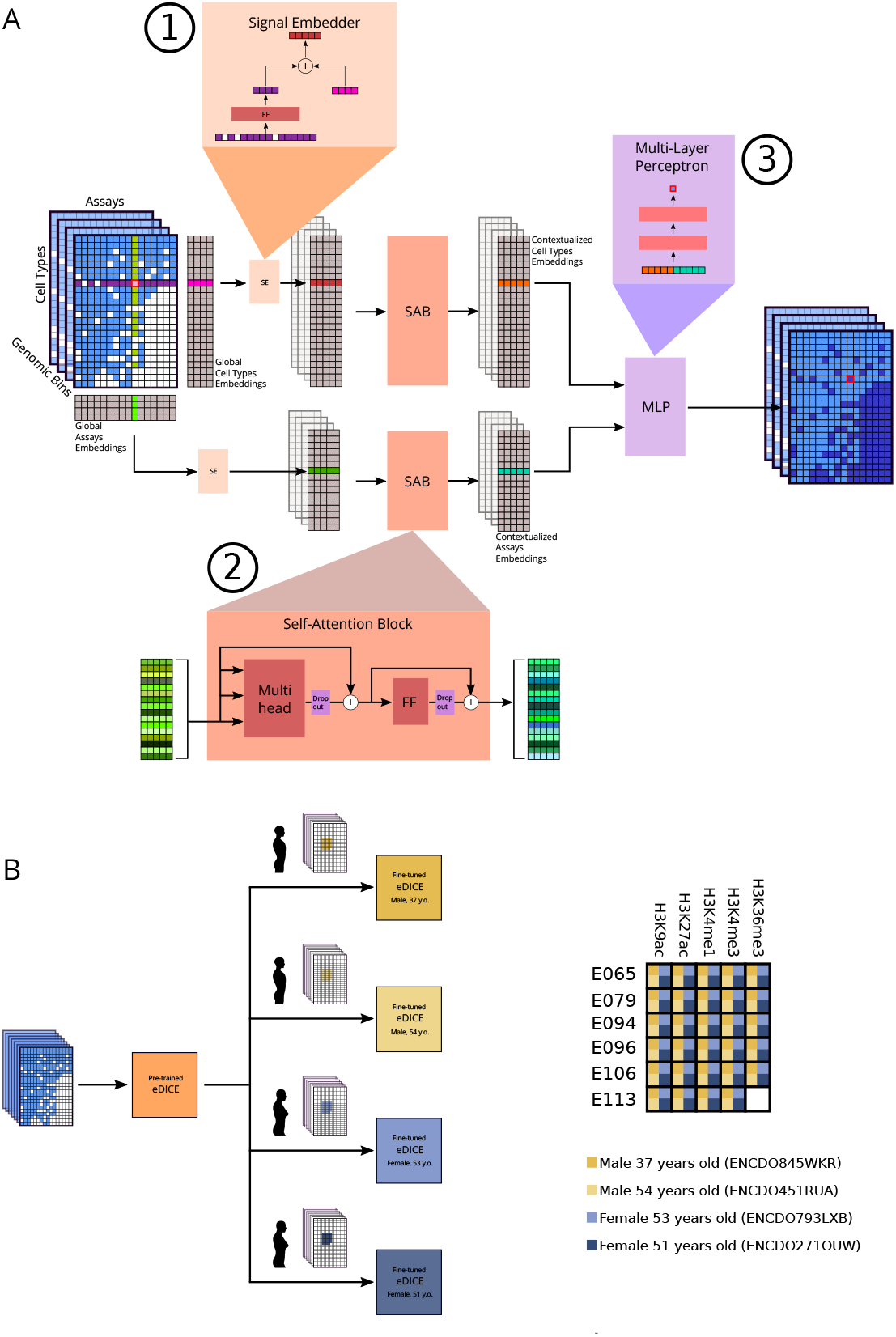
Schematic representation of the eDICE model. For each cell type we collect all measured signal values from assays performed in that cell type at the target bin, and project this set of values into a shared embedding space, where it is combined with a global embedding representing the cell type (1). We do likewise for assays, projecting the sets of values measured in different cell types from each assay into a distinct embedding space. We then apply self-attention over both sets of embeddings, allowing the network to capture relationships between cell types and assays to produce ‘contextualised’ embeddings which are functions of the local signal values in all observed tracks (2). Finally, a feed-forward neural network combines the contextual embeddings for a target cell type-assay combination to generate a prediction for the local signal value (3). **(b)** On the left, a transfer learning scheme used for the imputation of personalised epigenomes. On the right, the data matrix of the epigenomic tracks selected for the ENTEx case study is shown. It includes 29 tissue-assay combinations shared by all four individuals (for a total of 116 tracks).

To train the model, we simulate the imputation problem it will encounter at test time by randomly masking out a fixed fraction of the training signal values and using the remaining observed values to predict these simulated targets. This type of self-supervised task has been shown to be a meaningful training method for computer vision models[21]. This procedure is also reminiscent of few-shot regression approaches [22], where sets of observed input-target pairs sampled from a training distribution of regression tasks are used to learn models capable of producing task-conditioned predictions.

### eDICE imputations are highly accurate on the reference epigenomes

For direct comparison with previous imputation work, we evaluate the accuracy of eDICE imputations on a dataset of epigenomic tracks collated by the Roadmap project [23] and used in previous studies [13, 17, 18]. This dataset consists of 1014 signal tracks from 24 epigenomic assays in 127 cell types. All but one of the assays target histone modifications, with the remaining assay profiles chromatin accessibility via DNase-seq. A core set of five assays (H3K4me1, H3K4me3, H3K36me3, H3K27me3 and H3K9me3) is available in most cell types, while coverage of the cell types with the remaining assays varies widely. We use the first train/test split defined by [17], which consists of 709 training tracks, 102 validation tracks, and 203 test tracks. To compare the performance of imputation methods, we report a series of metrics assessing the quality of imputations of the tracks in the test set by models trained on tracks in the training and validation sets (and optionally using these tracks to provide inputs at test time). The metrics are computed across chromosome 21 of the hg19 assembly, the smallest human chromosome spanning about 48 million base pairs. As baselines, we report results for the prior methods ChromImpute and PREDICTD. We do not report metrics for Avocado since the publicly available imputations for the test tracks result from a model trained on the whole set of Roadmap tracks. However, Avocado’s performance on a similar set of metrics on the same dataset was reported to be very close to that of PREDICTD, with differences in performance tending to favour Avocado on MSE metrics and PREDICTD on measures of correlation and identification of peaks [18]. Finally, as a standard baseline, we also report predictions made by averaging the signal of the target assay in all other cell types except the target cell type (AVG).

Previous studies of imputation methods have varied in the choice of the primary metrics by which to assess performance [13, 17, 18]. In an attempt to provide a balanced view of model quality, we report performance on a selection of metrics designed to capture three desirable characteristics of imputations: (i) global similarity between imputations and ground truth values, (ii) similarity between imputations and ground truth values focusing on relevant subsets of the signal (e.g. high activity regions) and (iii) discrimination accuracy for a peak vs non-peak classification task. Metrics targeting the first category include the mean squared error and the Pearson correlation coefficient computed on the *arcsinh*-transformed signal for the whole genome. For the second category, we distinguish between foreground (Fg) and background (Bg) bins, where Fg bins correspond to the enrichment peaks detected by MACS2 [24]), and Bg bins correspond to the remaining bins, the complement to the fore-ground. For the third category, we rely on two different strategies for treating the continuous imputations as the outputs of a classifier. First, to test the ability of models to retain the semantic significance of the transformed p-value signal tracks, we use MACS2 as peak caller to detect the predicted peaks and compare them to the peaksets in the Roadmap dataset via precision and recall. Second, we use the threshold-agnostic area under the precision-recall curve (AUPRC) to offer a balanced view of the overall tendency of the models to produce imputations under which ground truth peak regions are ranked higher than non-peak regions, irrespective of the absolute predicted values. Additional details on all metrics are found in Supplementary Section S3.2.

The performance of the models is presented in Figure 2a and Supplementary Table S1. eDICE outperforms PREDICTD across all metrics, and ChromImpute across the majority, although ChromImpute shows strong performance for the prediction of peak height in the foreground. eDICE’s relative disadvantage here suggests a tendency to systematically underestimate the absolute signal values within peaks, which is exemplified in the trade-off between precision and recall compared to ChromImpute. However, it ranks peak and non-peak regions relative to each other more accurately than ChromImpute, as demonstrated by the fact that it outperforms all baselines on the AUPRC metric, thereby offering the best overall imputation in terms of global discriminatory power. We emphasise that while PREDICTD imputations were generated respecting the same data split, the ChromImpute imputations were produced in a leave-one-out fashion, so our model’s improved performance comes despite a considerable handicap relative to ChromImpute in terms of the available training data. Qualitatively, eDICE presents many of the characteristics that were present in its predecessors, such as a general smoothing of the imputed tracks, which is especially notable in the background regions (Figure 2a). Additionally, the imputed tracks reduce the impact of outlier values, such as the extremely high peaks present in a few tracks for H3K4me3. Such peaks are not necessarily a direct representation of the high significance of the local enrichment but can be heavily affected by the control samples’ coverage and quality, which, when low, can bias the estimated p-values towards extreme values.

**Fig. 2.**
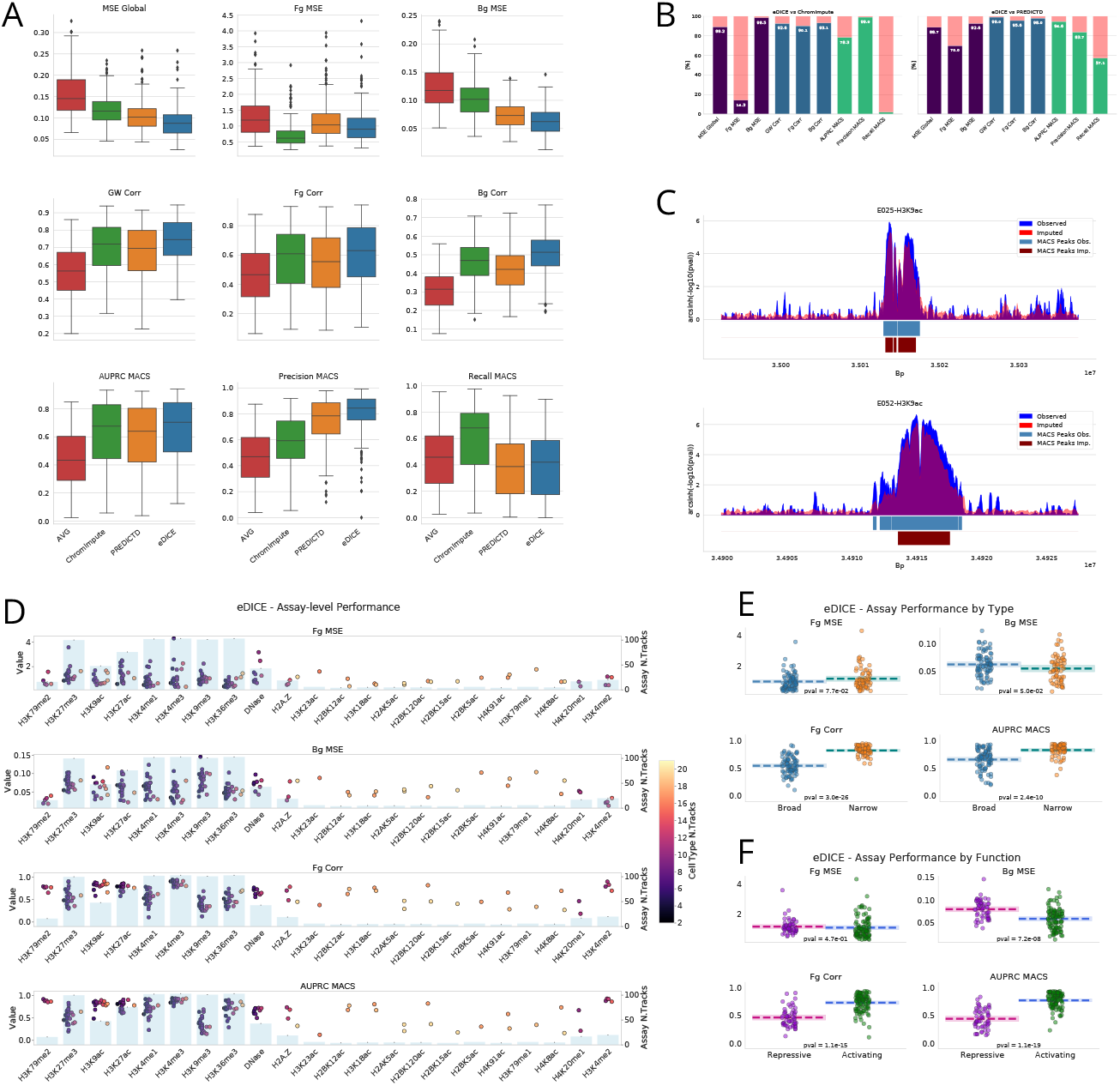
**(a)** Performance metrics for the imputation of the 203 test tracks on chromosome 21 for each model. **(b)** Percentages of test tracks on which eDICE outperforms the baselines for each metric. ChromImpute shows good performance on tasks related to the height of the peaks, while eDICE outperforms PREDICTD on all metrics. **(c)** Examples of observed epigenomic tracks with the signals imputed by eDICE. Below the tracks, the peaks detected with MACS2 highlight how the imputations accurately capture enriched regions. **(d)** Grouping the tracks by assay reveals significant differences in the imputation performance. This phenomenon is observed in the previous models as well, indicating that is most likely due to the nature of the specific modifications and the biases that their signal includes. The colour of each dot indicates the number of support tracks that share the cell type with that specific test track, while the light blue bars in the background show the number of support tracks that share the same assay. **(e)** Assays split into broad- and narrow-peak marks show consistently different performance for the imputation task. For each metric, we performed a 2-sided Welch’s t-test under the null hypothesis that both sets of metrics have the same mean, and reported the resulting p-value at the bottom of each plot. **(f)** Splitting the histone marks by functionality (repressive vs. activating) shows a similar bias as the comparison in (d).

To confirm that these aggregate results were not unduly influenced by variation in the range of metric values across different types of assay, we also examined the metrics at the level of individual tracks (Figures 2b and Supplementary Figures S4, S5) and aggregated by assay (Figure 2d). The track-level comparisons confirm that eDICE’s performance improvements are consistent across cell type-assay combinations. However, grouping tracks by assay reveals significant differences in the performance depending on the type of epigenetic mark. For example, all models tend to perform relatively poorly when predicting H3K27me3 and H3K9me3 (Figure 2c and Supplementary Figures S6 and S7). Comparing the average assay-level performance of each model shows that the improvements brought by eDICE are consistent across the board despite these discrepancies between assays (Supplementary Figure S2).

Finally, we further explored whether differences in performance between types of assays could be related to differences in specific properties of the epigenetic marks. Some histone modifications can be classified as either narrow-peak (H3K27ac, H3K4me2, H3K4me3, H3K9ac) or broad-peak marks (H3K27me3, H3K36me3, H3K4me1, H3K79me2, H4K20me1). Comparing the performance of eDICE on test tracks across these two groups, we observed that performance tended to be higher on narrow-peak than on broad-peak marks for correlation and classification metrics (Figure 2d). Furthermore, a similar divide is observed when splitting histone modifications into repressive (H3K27me3, H3K9me3) and activating marks (the active promoter-associated H3K9ac, H3K4me2, H3K4me3, active enhancer associated H3K4me1 and H3K27ac and DNAse-seq, Figure 2e). As repressive marks are often linked to heterochromatin configurations, this discrepancy is possibly due to biases introduced by the processing pipelines because of systematic sequencing differences in these regions. However, as repressive marks also tend to display broad peaks, it is challenging to pinpoint the precise reason for the observed differences.

### Imputations accurately capture significant differences between tissues

Epigenomic patterns differ between cell types to control and register cell function and identity. It is critical that imputations accurately capture these differences if they are to constitute valuable resources of cell-type-specific epigenomic landscapes. However, within the scale of the whole genome, these cell-type-specific differences are subtle and global evaluation metrics such as those considered above are dominated by regions which have a shared functionality across cell types, such as large intergenic regions.

We next seek to evaluate the usefulness of our imputations in identifying cell-type-specific variability. As a test case, we identify differential peaks across two imputed H3K9ac tracks from the test set (corresponding to Roadmap cell types Adipose-Derived Mesenchymal Stem Cell Cultured Cells (E025) and Muscle Satellite Cultured Cells (E052)). Notably, experimental protocols to identify differential patterns explicitly require several biological replicates to estimate local variability, which is essential for robust statistical hypothesis testing [25] (Figures 3a and 3b). Typical imputation methods, however, pool biological replicates and focus on predicting the mean signal (bottom parts of Figures 3a and 3b).

**Fig. 3.**
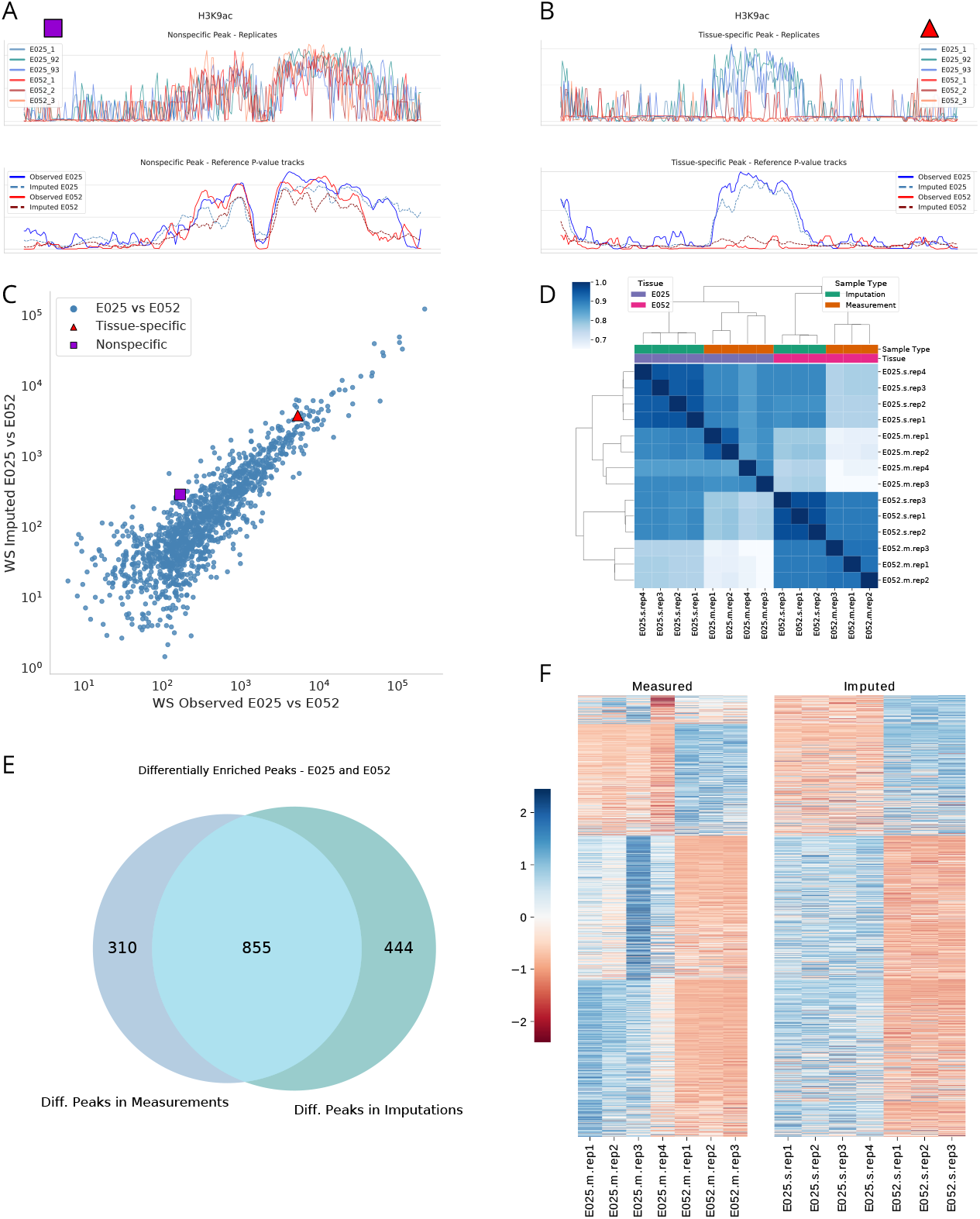
Differential peak analysis using imputed epigenomic tracks. **(a)** and **(b)** show examples of nonspecific and tissue-specific peaks respectively for H3K9ac in the two chosen tissues (E025 and E052). The upper part shows the measured replicates, emphasising the need to account for the biological variability in future improvements of epigenomic imputation models. **(c)** A scatterplot of the Wasserstein distance between the signal in the two tissues, for each peak in the enriched peakset of E025. The x axis displays the WS distance between observed signals, while the y axis between imputed signals. The imputations retrieve most of the information contained in the measurements, especially for the stronger differences between tissues. We highlighted the two points corresponding to the peaks shown in figures and (b). **(d)** Correlation heatmap of the affinity scores for different replicates. The simulated replicates correctly retrieve the expected relationships to the measured replicates, although they show a high degree of similarity between themselves, likely an artefact of the simulation procedure. **(e)** Venn diagram representing the peaks that are detected as differentially enriched between tissues using imputed and measured signals. The imputed signal retrieves 66% of the true peaks. **(f)** Binding affinity heatmaps for the measured replicates and the imputed pseudo-replicates. Each row corresponds to one of 1609 differentially enriched peaks detected in either of measurements and imputations. The imputed replicates display the same global block structure as the measurement replicates.

We first note that the overall shape of individual peaks is remarkably conserved between individual experimental replicates for corresponding tracks (Figure 3a). In the case of tissue-specific peaks, on the other hand, the signal shapes are distinct between replicated measurements derived from different tissues (Figure3b). We have previously exploited this observation for differential peak calling [26], where we considered the genomic region of the peak as a metric space and treated the pile-up of sequenced reads like a sample from a hidden probability distribution on that space. This strategy dramatically improves the test’s statistical power compared to methods based on total counts alone. We also note that the shape differences are well captured by the mean signals (Figures 3a and 3b bottom panels). To quantify differences in peak shapes across the two cell types, we computed the Wasserstein (WS) distance between the pooled ground truth signals across the two cell types, and likewise between the imputed signals. Figure 3c shows that the distances in imputed and ground truth tracks are strongly correlated, indicating that the imputations accurately capture cell-type-specific differences in the shape of signal enrichment at peak regions.

As an independent analysis, we next took advantage of the robustness of the existing differential peak analysis method, DiffBind [27], which, however, requires replicates for statistical testing. We have therefore estimated the local variability of cell-type-specific test tracks. Assuming a negative binomial distribution, the estimated variance parameters are subsequently used to simulate replicates from the imputed mean signal tracks on chromosome 21. While an arbitrary number of replicates can readily be generated in this way, we chose to use three simulated replicates, similar to typical experimental scenarios. Those tracks were fed into the standard differential analysis pipeline, and the outcome is compared with the results obtained from the corresponding analysis of actual replicated measurements. Further details on the simulation procedure are provided in Supplementary Section S3.4. We emphasize that the simulation procedure employs only replicates from the training set and tissue-specific control samples in addition to the imputed tracks and makes no use of any information from the test set.

Employing the DiffBind library [27] we compare *binding affinity scores*, which are indicative of the strength of interaction between DNA and biomolecules (such as modified histones). Figure 3d shows a correlation heatmap for the similarity of affinity scores for different samples. The block structure highlights the expected relationship between the replicates derived from different tissues; however, the simulated replicates show high similarity across tissues, possibly due to the adopted procedure underestimating the biological variance between samples.

Within DiffBind, we used DESeq2 to identify peaks of differential enrichment with default parameters. Specifically, we used a ‘local’ fittype to estimate dispersion and used a Wald test for negative binomial distribution (‘nbinomWaldTest’) to identify statistical significant peaks. A total of 1165 and 1299 peaks were detected as differentially enriched in measurement and imputations, respectively (FDR threshold of 0.05). 855 peaks (~ 73% of the measured peaks) are shared between the two sets, resulting in a Positive Predictive Value of 0.66 (Figure 3e).

Details of the binding affinity scores for each differentially enriched peak in the consensus peakset derived from imputations and measurements are shown in Figure 3F, where the block structure of the measurements (left side) is replicated in the imputations (right side).

In summary, we conclude that the imputations accurately capture cell-type-specific differences, both in terms of altered shapes of signal enrichment at peak regions and also with regard to integrated total counts in the peak regions, when considering local variability.

### eDICE accurately predicts global histone modification patterns from individual donors

Recently, a collaboration between the ENCODE [16, 28] and the Genotype-Tissue Expression (GTEx) consortia created new data sets that include extensive individual-specific histone modification measurements from four donors. This constitutes an important resource and a significant step towards personalized functional genomics. However, the large-scale measurement of comprehensive epigenomic maps for individual healthy donors or patients continues to be prohibitively expensive and imputation methods, like eDICE, are opening up the possibility of expanding personalised epigenomic maps using only a small set of assayed marks.

The ENTEx data set includes data across 25 different tissues from two adult males, 37 and 54 years old, and two adult females, 53 and 51 years old. To test eDICE’s capability to impute individual-specific tracks we have selected 29 tissue-assay combinations available for all four individuals (Figure 1b). These tracks are chosen so that the histone marks and tissues approximately overlap with those present in the Roadmap dataset. We extract data from chromosome 21 to be used in the following experiments. Additional details on the selected sets can be found in Supplementary Section S3.5, together with the accession codes for the ENTEx experiments (Supplementary Table S2).

We first note that many individual-specific tracks are largely similar across the four different individuals, as seen for instance for the H3K4me3 tracks in E106 (Figure 4a). We also find that cell identity is a dominant determinant of epigenomic patterns, in particular for marks H3K27ac, H3K4me1 and H3K9me3 (Figure 4b). However, there are also individual-specific epigenomic signatures, most notable for H3K9me3, with local enrichment unique to only one or a subset of individuals (Figure 4c). These personal epigenomic differences may either reflect underlying DNA sequence variants, in which case they may be observable across different tissues of the same individual, or they may result as a consequence of ageing or due to interactions with external stimuli, potentially in a tissue-specific manner. Three-dimensional histograms of co-occurrences across tissues and individuals show striking differences between histone marks (Figure 4d and Supplementary Figures S15-S19); overall we find indivdual-specific enrichments that are shared across all examined tissues to be rare. However, there is a large number of individual-specific peaks that are uniquely found in one tissue, particularly for mark H3K9me3. While individual epigenomic signatures are rare in the whole genome-wide context, they may be highly informative, and could be used for personalised predictions for risk stratification [29], drug resistance [30, 31], or personalised therapies [32]. Therefore, when imputing epigenomic tracks in an individual-specific manner we want to accurately recapitulate the overall cell-type signatures, while at the same time capturing the subtle individual differences.

**Fig. 4.**
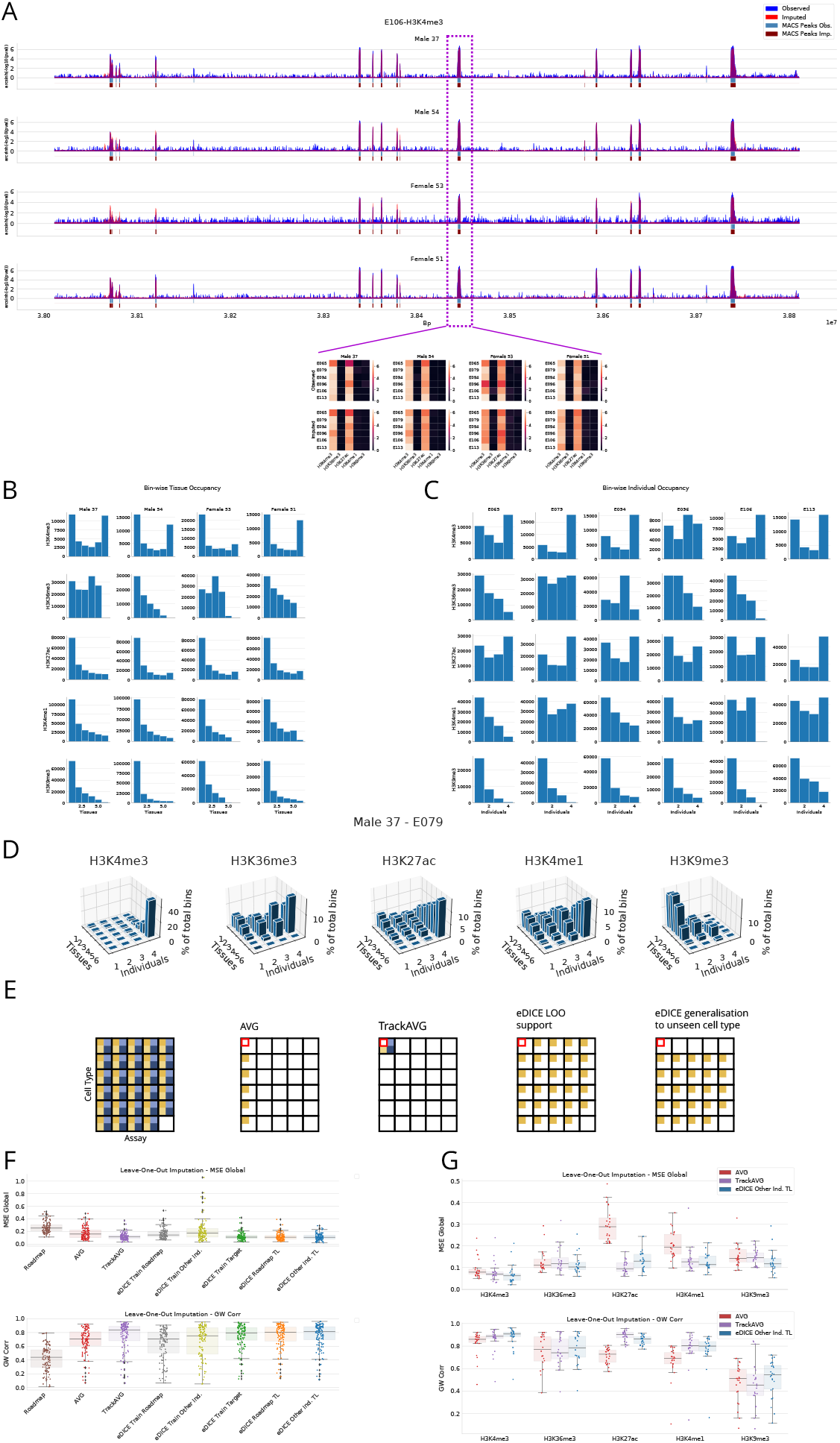
**(a)** E106-H3K4me3 track spanning 800kb and showing consistent patterns for all four individuals. For the central peak we display a slice across the epigenomic tensor demonstrating signal conservation across tissues and individuals. **(b)** Occupancy histograms for the enriched bins across tissues. **c)** Occupancy histograms for the overlap of enriched bins across individuals. **(d)** Occupancy across tissues and individuals for each enriched bins in the tracks for Male 37 for the E079 tissue. **(e)** Data and configurations used for the baselines and the eDICE models. AVG averages the respective assay tracks over tissues from a given individual. TrackAVG averages cell-specific tracks from different individuals. The latter 2 plots highlight the leave-one out and the out-of tissue strategy used to train and test eDICE models. **(f)** MSE and Pearson correlation for the LOO imputation of chromosome 21 using a variety of training schemes. TrackAVG demonstrates good genome-wide performance. Among the eDICE setups, the best performing model is trained on 3 individuals and tuned to the target individual. **(g)** Performance metrics split by assay for the task in (e). The imputations show large variations between the various histone modifications.

We first examine the global similarity between imputations and measured tracks using MSE and Pearson correlation. Subsequently, we focus on the detection of individual-specific signatures, quantifying performance using area under the precision-recall curve (AUPRC).

As baseline methods, we include several imputation-free strategies. Firstly, we consider the signals in the corresponding tracks in the Roadmap data set; secondly, we average signal values in the target cell-type-assay track across the three non-target individuals (TrackAVG, see Figure 4e). By design, these baselines reflect the characteristic signatures of the target assay and tissue, but not the individual-specific enrichment that characterises each person. As a third baseline, we take the average value of the target assay in all other cell types for which it is available in the chosen individual (AVG) (Figure 4d). This last strategy is able to capture individual-specific signatures, however, it is not cell-type specific.

We next explore several training strategies for eDICE. In each case we assume a leave-one-out (LOO) setting where we have 28 tracks available from the target individual and wish to predict the remaining track (Figure 4d)). Three models are trained from scratch on the complete Roadmap dataset (eDICE Train Roadmap), the target individual only (eDICE Train Target), or the three off-target individuals (eDICE Train Other Ind.), to then use the 28 tracks from the target individual only as input. Two models adopt a transfer learning (TL) approach where they are first trained on either the complete Roadmap dataset (eDICE Roadmap TL) or the three off-target individuals (eDICE Other Ind. TL), and then fine-tuned using the 28 available tracks before predicting the target track. For both TL configurations, we tune the signal embedder and the final MLP while freezing the global assay and cell type embeddings and the transformer block, with the aim of preserving as much information about the relationships between assays and cell types as possible.

Figure 4f shows the MSE and Pearson correlation for the LOO imputation of the 29 ENTEx tracks on chromosome 21.

We first note how the TrackAVG baseline performs remarkably well for this task, indicating that global cell-type-specific enrichment patterns are highly conserved across individuals. Several configurations of eDICE offer competitive performance, specifically the transfer learning approaches. On the other hand, the models that do not use any data from the target individual for training or fine tuning (eDICE Train Roadmap and eDICE Train other Ind.), display sub-optimal performance, highlighting the need to account for the individual-specific biases. These may include genuine individual-specific patterns, as well as systematic errors (such as batch effects) that may affect the measurement for that specific individual.

Another advantage that the TL eDICE models offer compared to models trained from scratch is a considerably reduced computational cost for training. The fine tuning or transfer learning process is significantly faster than the training process, and optimises a smaller number of parameters. For example, the eDICE Train Target model required 50 epochs of training for optimal performance, while the eDICE Other Ind. TL model achieved better performance with only 5 epochs of tuning.

Among the eDICE configurations, the transfer from other individuals offered the best results, both in terms of performance metrics and computational costs, which suggests that personalised imputation models are potentially impacted by several sources of bias. Previous applications of deep learning to epigenomic measurements have highlighted how model performance is significantly affected by distributional shifts [33]. In this case, transfers between datasets need to account for large changes in experimental conditions (evidenced in the models trained on Roadmap) but even within the same dataset individual-specific biases are significant (clearly shown in the performance of eDICE Train Other Ind.).

Analysing the performance at the level of individual assays (Figure 4G) reveals significant differences between histone modifications. The H3K9me3 assays, for example, are the most difficult to predict as they exhibit many individual-specific as well as tissue-specific patterns (Figure 4 D). However, on this mark, the eDICE transfer learning strategy is particularly effective, showing improved performance relative to the baselines.

### eDICE captures epigenetic variation between individuals

Defining individual-specific epigenomic signatures is far from trivial; a robust analysis would require more than four individuals to properly understand the overlap of enriched regions and what external factors influence it. Here, we define individual-specific signatures as the set of enriched regions detected from the measured samples that span at least 150bp and which are present only in one individual. This definition aims to capture peaks such as the example shown in Figure 5a, where H3K4me3 is clearly found in one individual but not the others. This task presents significant challenges due to the small portion of epigenetic enrichments that meaningfully differ between individuals and because of the complex epigenetic patterns that arise in these regions of variability, exemplified by the heatmaps displayed in the lower portion of Figure 5a). In these cases, local variability is observed not just between individuals but also between tissues within the same individual (Figure 5b).

**Fig. 5.**
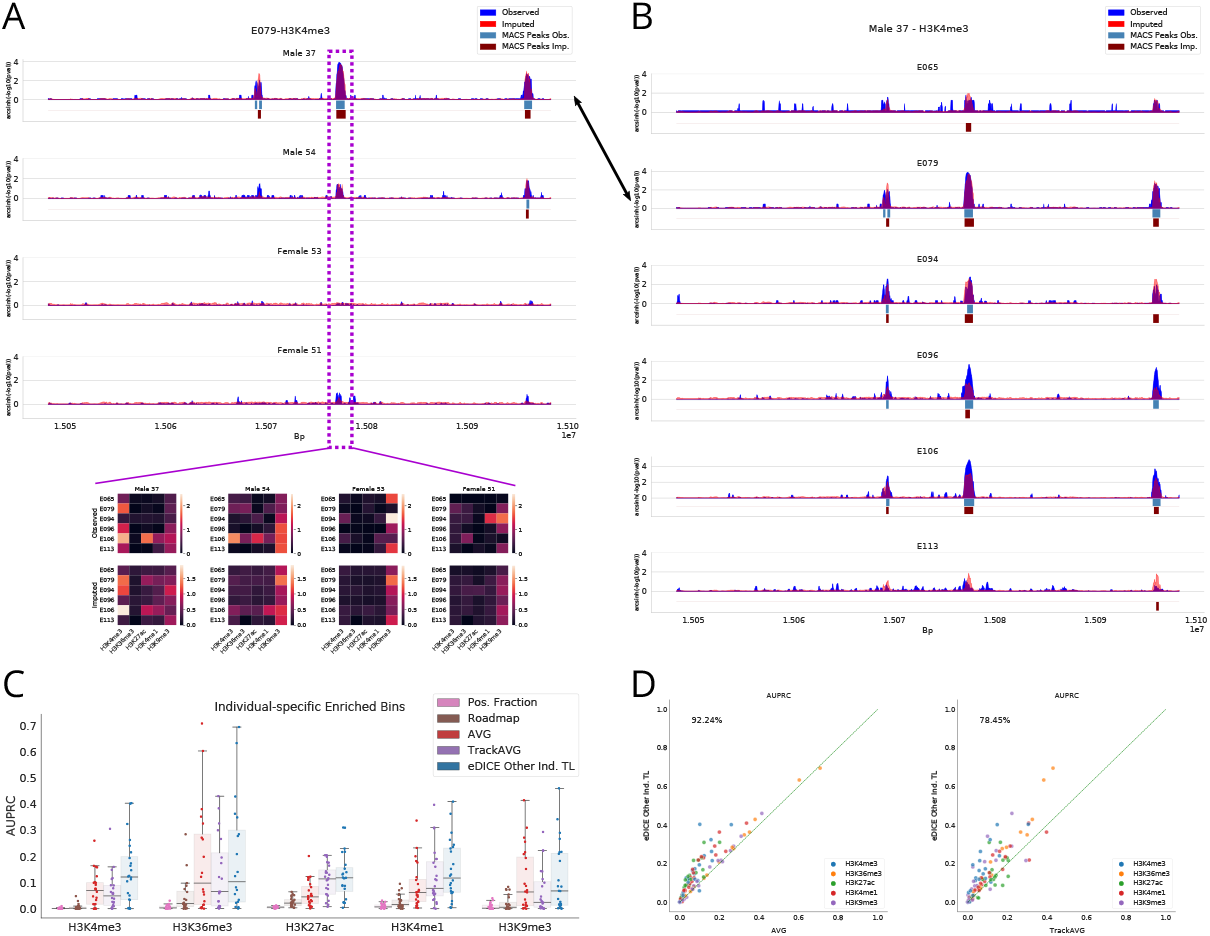
**(a)** Individual-specific H3K4me3 enrichment for (Male 37) in tissue E079. For this genomic location, we display a slice of the epigenomic tensor for each of the four individuals, highlighting the challenge of imputing these varied patterns. **(b)** Observed and imputed tracks for the H3K4me3 assay in Male 37 across tissues in the same genomic region as (a). **(c)** AUPRC for the prediction of individual-specific enrichment in the LOO ENTEx imputations, where the peaks shared with other individuals have been masked out. **(d)** Track-level AUPRC for the prediction of individual-specific enriched bins. eDICE improves on the AVG baseline on almost 90% and on 78% of the tracks when compared to TrackAVG.

We mask out the enriched regions shared between individuals and measure the capability of imputation models to correctly classify the individual-specific signal using the area under the precision-recall curve (AUPRC). Here we test the best eDICE configuration from the previous experiment (eDICE Other Ind. TL). For completeness, we included the fraction of positive samples (Pos. Fraction) as a reference baseline for the AUPRC measure [34]. The results, presented in Figure 5c, show that eDICE considerably improves the prediction of individual-specific enrichment compared to the AVG and TrackAVG baselines. A track-level comparison of eDICE’s improvement over AVG is shown in Figure 5d.

### eDICE generalises to unseen tissues through transfer learning

The transfer learning procedure allows us to test the imputation performance of eDICE when imputing epigenomic marks for cell types that were not observed for the target individual. We extend the LOO procedure to exclude all tracks for a specific tissue at one time, which are then imputed using the remaining data (last configuration presented in Figure 4e).

We test both TL configurations, generalising from the Roadmap dataset and from the off-target individuals, against the AVG and TrackAVG baselines unaltered from the previous experiments.

Figure 6a displays the MSE and Pearson correlation for the imputation of the 116 ENTEx tracks with the previously described procedure. Notably, the Roadmap data is a pure approximation of the respective measurements from individual donors. Again, averaging strategies either across tissues of the same individual or within the same cell type across different individuals result in tracks that are overall very similar to the actual measurement, however at the cost of loosing individual- or tissue-specificity. The transfer learning strategy from the Roadmap dataset still offers some improvements over the baselines, which opens up prospects for generalising to a large set of tissues when mapped in large-scale efforts such as Roadmap. Consistent with expectations, the transfer between individuals outperforms the Roadmap TL (Figure 6b), most likely due to the shared experimental conditions within the same dataset that require a much less extensive adaptation in the fine-tuning of the model when compared to changes between datasets.

**Fig. 6.**
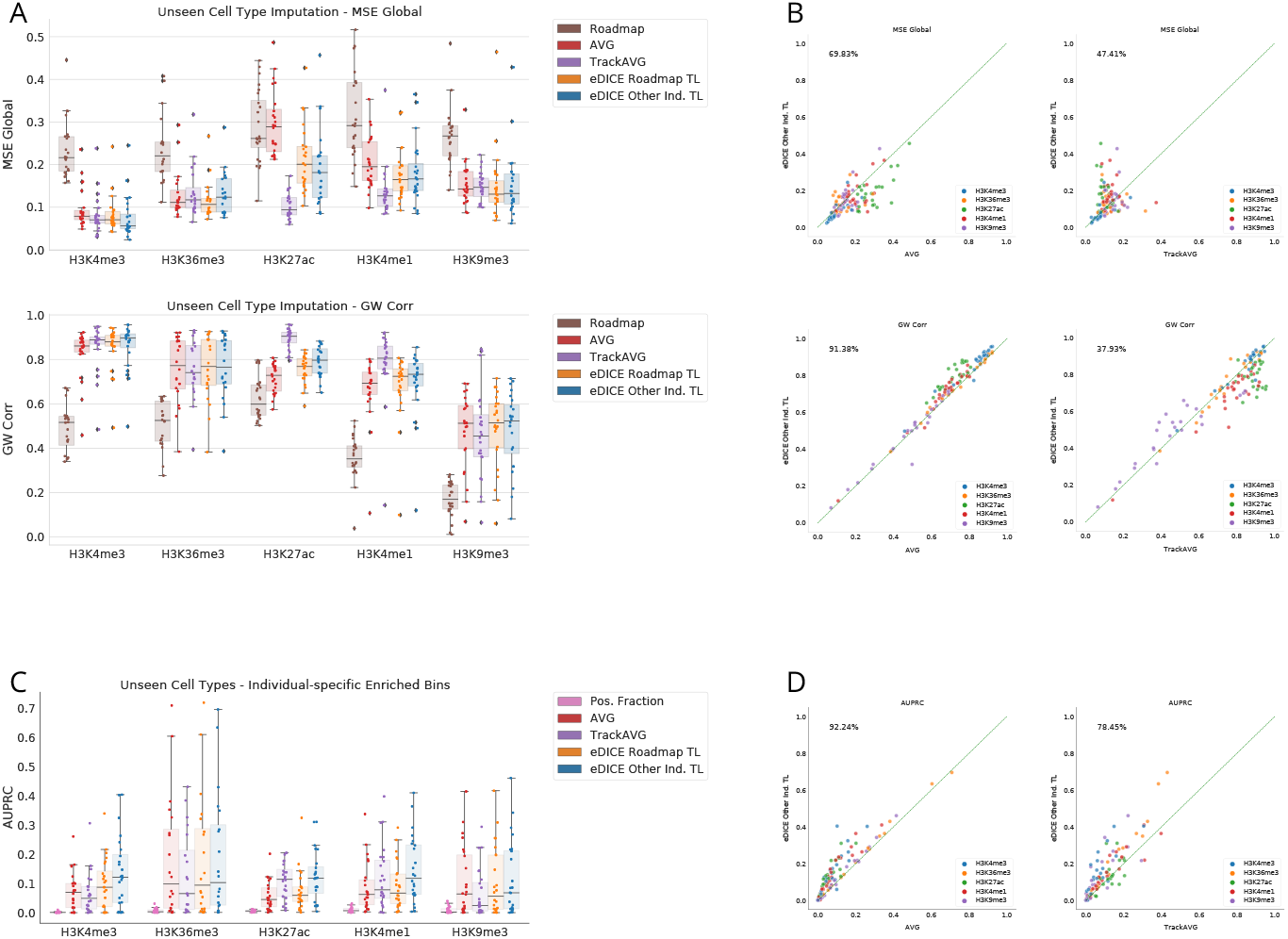
**(a)** MSE and Pearson correlation for the imputation of tracks for unseen cell types. Transfer learning between individuals offers the best performance, but remarkably a model trained on the Roadmap dataset and tuned to the target individual still offers improvements compared to the AVG baseline. **(b)** Track-level comparison of the imputation metrics for the generalisation to unseen cell types, highlighting the percentage of tracks in which eDICE outperforms each model. **(c)** AUPRC for the prediction of individual-specific enrichment in the imputed tracks for unseen cell types. **(d)** Comparison of the AUPRC for the predictions of individual-specific enrichment in (c). Transfer learning between individuals clearly outperforms the AVG baseline and the transfer learning from the Roadmap dataset.

Once again, clear differences between histone marks emerge from these plots, suggesting that the use of imputation models might be more suitable for certain epigenetic marks than others.

Following the same setup for the prediction of individual-specific enrichment, we observe that eDICE retains a small advantage over the AVG baseline when generalising to unseen tissues (Figures 6c and 6d).

This result highlights that imputation models offer the possibility of capturing personalised epigenomic signatures even in more restrictive conditions, which is a promising direction for the application of epigenomic imputations for precision medicine.

## Discussion

We have presented eDICE, a novel epigenomic imputation framework that outperforms state-of the art methods on several metrics, while combining the advantages of its predecessor models. Like ChromImpute, eDICE uses the local signal of observed tracks to encode information on the genomic position, removing the need to learn explicit embeddings for each genomic position. Similar to the tensor factorisation models PREDICTD and Avocado, eDICE uses factorised representations for computational efficiency, and readily generalizes to cell type-assy combinations unseen during training, while at the same time requiring a three orders of magnitude lower number of parameters than Avocado.

The choice of evaluation metrics for epigenomic models is far from trivial. We selected a wide array of metrics that capture information relevant to possible downstream tasks in which epigenomic imputations would be used; on most metrics eDICE outperforms all baselines, with the exception of a few where ChromImpute shows great performance.

We emphasize the need for imputation models to be trained and designed with the aim of including imputations in established bioinformatics processes. As a case study, we explored the possibility of simulating biological replicates from the imputed data, which are then used for differential peak calling obtaining results compatible to the measured replicates. We pose that future developments in the field of epigenomic imputation should account for and predict not only the average value of measurements, but also the intrinsic biological variability of different samples. Explicitly modelling the variance of epigenomic measurements would allow for more robust analysis to distinguish the differences caused by fluctuations due to the natural variability of the samples from the true differences between tissues and marks that encode the functional variations of cell profiles.

Finally, we demonstrated the possibility of imputing personalized epigenomic tracks with eDICE. Simple baselines obtain competitive performance in capturing the global patterns shared across tissues and individuals. However, this case study shows that eDICE is better able to predict the individual-specific enrichment that is not captured in those baselines. While still a proof of concept, the transfer learning framework adopted allowed eDICE to generalise to unseen cell types while still retaining improved prediction of individual-specific enrichment. The extremely small number of individuals in the ENTEx dataset is a severe constraint for a robust study of machine-learning-assisted personalised epigenomics. Nevertheless, imputation models such as eDICE open up exciting opportunities for future precision medicine workflows, in which a small set of measurements from a patient would be used to obtain a more complete epigenomic map.

Future developments of eDICE and other imputation methods should include explicit modelling of the natural variance present in biological replicates. Additionally, much work remains to be done to understand the systematic biases introduced by bioinformatics pipelines used to process sequencing data; it is possible that a shift from p-value signals to read counts could improve the robustness of the methods and reduce the noise caused by shallow control replicates. We suggest that in-depth analysis of the shifts between datasets is key for the practical applications of epigenomic imputations. Supplementing the limited information contained in small experiments with the large scale maps gathered by international consortia offers a promising methodology to overcome the experimental constraints that limit our understanding of epigenetic marks.

## Online Methods

### Data

The dataset chosen to evaluate eDICE is the set of epigenomic measurements from the Roadmap Consortium [12]. The Roadmap dataset consists of 1014 signal tracks from 24 types of epigenomic assay in 127 cell types. All but one of the assays target histone modifications, with the remaining assay profiling chromatin accessibility via DNase-seq. A core set of five assays, targeting H3K4me1, H3K4me3, H3K36me3, H3K27me3 and H3K9me3, is available in each cell type, while coverage of the cell types with the remaining assays varies widely. We use the first train/test split defined by [17], which consists of 709 training tracks, 102 validation tracks, and 203 test tracks. Supplementary Figure S1 gives an overview of the data splits over train, validation and test.

These signal tracks are obtained by mapping a set of sequence reads to a genome to form a genome-wide activity profile. A signal track is identified by the assay used to generate reads and the cell type for which the assay was performed. Given a set of observed tracks that are the result of performing at least one of a set of *n*_*a*_ assays 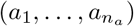 in each of a set of *n*_*c*_ cell types 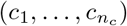, the goal is to generate imputations for all assay-cell type combinations which are not represented by tracks in the observed set.

Following previous imputation work, we work with signals in the form of − log 10 p-value tracks, which indicate the statistical significance of a mark at each genomic position, and seek to impute the average − log 10 p-value within each non-overlapping 25 base pair interval in a given subset of the genome.

We additionally preprocess the log 10 p-value signal using an *arcsinh* transform, which reduces the impact of outliers and differences in distribution between different types of assay, again inspired by prior work [17, 18, 35].

The 116 tracks selected from the ENTEx dataset (Supplementary Table S2) have been processed in the same manner.

### Factorised representation learning

The imputation problem can be seen as one of combinatorial generalisation in which, given a set of combinations of cell types and assays for which signal tracks are fully observed, we predict full signal tracks for unobserved combinations of cell types and assays at test time. Such problems are naturally solved using a factorised approach, in which predictions are generated as a function modelling the interaction between separate representations for each element of the combination.

One such method is the tensor factorisation framework adopted by PRE-DICTD and Avocado, in which the factorisation is extended to the genomic position axis. Learned cell type embeddings, **c**, assay type embeddings, **a**, and bin embeddings, **b**, are combined via a parametric function *g*_*θ*_ to reconstruct or impute tensor elements:

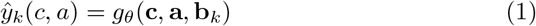

We follow the use of a factorised structure for representing cells and assays, but instead of learning independent embeddings for each bin, we allow the observed signal values from all tracks at a given genomic position to condition the cell and assay representations at that position. Let 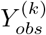 represent the set of signal values in all observed tracks at the *k*^*th*^ bin. We propose a (local) factorisation of the form:

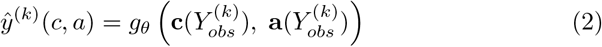

in which cell type and assay representations are independently conditioned by the local signal before being combined by an output network. We refer to the observed tracks in 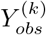 that are fed as input to the model as *support tracks*, and to tracks whose values are predicted as *target tracks*.

### eDICE model

The model produces the local embeddings 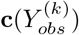 and 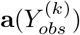 by mapping input vectors representing the local signal associated with each cell type and each assay to sets of contextualised local cell type and assay embeddings via separate Transformer-style self-attention block which act separately on cell types and assays. We describe the architecture below and schematise the main steps in Figure 1.

We first describe the construction of the inputs to the self-attention blocks. Let 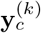 denote a partially observed signal vector characterising the signal in all assays in cell type *c* in the *k*th bin. Then

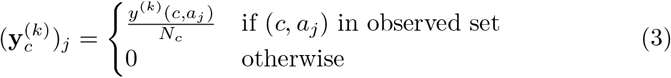

where *N*_*c*_ denotes the number of observed pairs (*c*, *a*_*j*_) in the local signal data, i.e. the number of non-zero elements in the vector 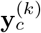. This scaling factor limits the changes in the norm of this local signal vector due to uneven mapping of the epigenome, and is equivalent to the activation scaling used in Dropout [36].

This cell-specific local signal vector is mapped to an embedding space through a non-linear function 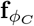, shared by all cell types, and implemented through a fully connected layer with parameters *ϕ*_*C*_ and a ReLU activation function. To allow the network to combine the local signal representation with knowledge of the global properties of the cell type, we add to the local signal embedding a learned global cell type embedding **u**_*c*_, which plays the role of a position embedding in the standard Transformer architecture.

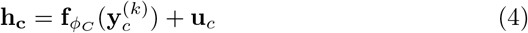

We note that in order to keep the notation simple, we are suppressing the bin superscript in the intermediate cell type embeddings **h**_*c*_. These embeddings are passed as input to a multiheaded self attention block (*SAB*)(•), which has the same structure as that used in the Set Transformer architecture except for the omission of layer normalisation.

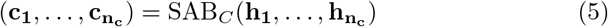

The same methodology is applied to produce the contextual embeddings for the assays **a**:

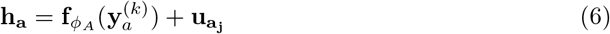

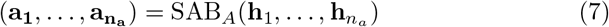

The result of the factorised self-attention is a set of cell representations 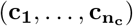 and a set of assay representations 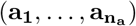, each of which is a function of the identity of the particular entity being represented and the full set of local signal values in all observed tracks (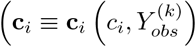 and 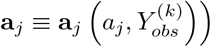) Given these representations, the

corresponding contextual cell type and assay representations through the fully-connected neural network *g*_*θ*_ (Equation 2).

### Hyperparameters and training details

The model uses cell and assay embeddings of dimension 256 at all stages in processing. Within the self-attention block we use 4 attention heads, that are then fed to a feed-forward neural network with a single hidden layer with 128 neurons and a 256-dimensional output. Finally, the combination of cell and assay representations are fed to a multilayer perceptron with 2 hidden layers with ReLU activations and 2048 neurons per layer. During training Dropout with rate 0.3 is applied to each hidden layer in the output MLP.

The hyperparameter configuration was selected based on the best performance on the validation set, and the final model was retrained on the combination of training and validation sets.

The model is trained to reconstruct the signal values for randomly selected subsets of the observed tracks, whose values are masked from the inputs. The number of masked tracks is constant. This training objective can be seen as a kind of self-supervised learning, similar to that employed by denoising autoencoders [37]. The model used to analyse the eDICE performance in the Results section was trained on the union of the training and validation set for 50 epochs, using the Adam optimiser with a learning rate of 3 * 10^−4^, and using 120 randomly selected tracks as targets for each training bin.

For the ENTEx imputations, the “eDICE Train Roadmap” model was trained with the same exact configuration just described but including all 1014 Roadmap tracks in the training set. “eDICE Roadmap TL” used the same model as starting point and further tuned the signal embedder and MLP for 20 epochs with a learning rate of 10^−4^. The best performing “eDICE Train Other Ind.” model used an embedding dimension of 64 and 512-dimensional hidden layers in the MLP and was trained for 20 epochs with a learning rate of 10^−4^. The “eDICE Other Ind. TL” used the same pre-trained model to tune for 5 epochs with a learning rate of 10^−5^. “eDICE Train Target” was trained for 50 epochs with a learning rate of 3 * 10^−4^.

## Supplementary information

Additional information for this paper is offered in the Supplementary Materials.

## Acknowledgments

The authors thank the International Max Planck Research School for Intelligent Systems (IMPRS-IS) for supporting Tanmayee Narendra.

## Declarations

### Funding

This project has received funding from the European Union’s Framework Programme for Research and Innovation Horizon 2020 (2014-2020) under the Marie Skłodowska-Curie Grant Agreement No. 813533-MSCA-ITN-2018. GS acknowledges funding from the Academy of Medical Sciences, Springboard and the UKRI Future Leader Fellowship. This work was supported by the BMBF-funded de.NBI Cloud within the German Network for Bioinformatics Infrastructure (de.NBI) (031A532B, 031A533A, 031A533B, 031A534A, 031A535A, 031A537A, 031A537B, 031A537C, 031A537D, 031A538A)

### Data availability

The Roadmap dataset is available at http://www.roadmapepigenomics.org/ The epigenomic tracks for the 4 individuals part of the ENTEx dataset can be found on the portal for the ENCODE project https://www.encodeproject.org/.

### Code availability

Source code for eDICE can be found at https://github.com/alex-hh/eDICE. The processed HDF5 files containing the training bins and chromosome 21 for the Roadmap dataset, and chromosome 21 for the selected tracks of the ENTEx dataset can be provided upon request.

### Authors’ contributions

AHH designed and implemented the model imp for the ENCODE imputation challenge as well as its subsequent improvements eDICE, contributed to the performance validation on the Roadmap dataset and to the writing of the manuscript. GV implemented and performed the analysis of the ENTEx dataset, contributed to the Roadmap validation, the differential peak analysis, the implementation of the transfer learning strategy and the writing of the manuscript. TN implemented the differential peak analysis and contributed to the writing of the manuscript. MR-C contributed to the supervision of the ENCODE Challenge contribution. BS has contributed to the conception of the work. GS supervised the development of the ENCODE imputation challenge method imp and eDICE, participated in the analysis of the Roadmap imputation, the differential peak analysis and the ENTEx imputations, and contributed to the writing of the manuscript.

## Notes

### Competing Interest Statement

The authors have declared no competing interest.

