## Supplementary Material for "Getting Personal with Epigenetics: Towards Machine-Learning-Assisted Precision Epigenomics"

### Supplementary Materials

#### S1 Supplementary Figures

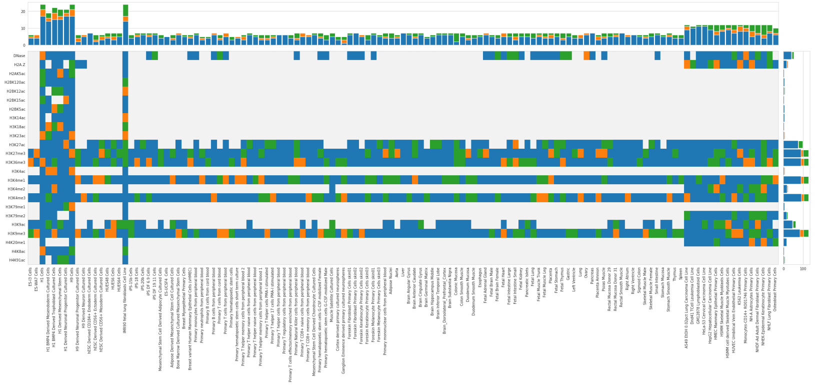


**Fig. S1** The data matrix showing which tracks from the Roadmap consortium were part of the training (blue), validation (orange) and test (green) sets.

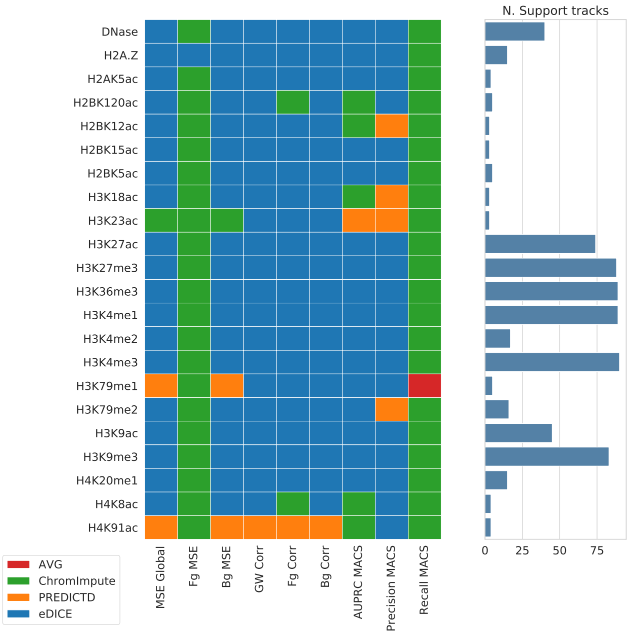


**Fig. S2** Heatmap that showcases which model presents the best average test performance for each assay, divided by metric.

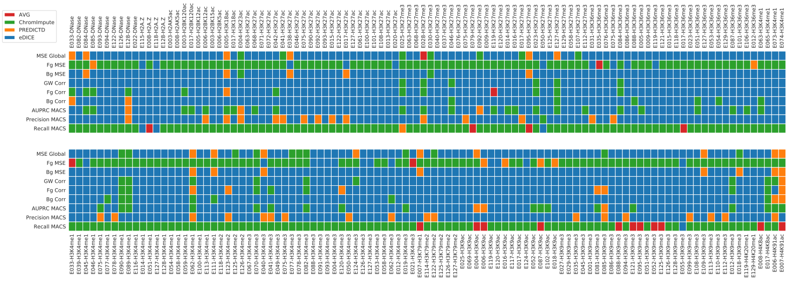


**Fig. S3** Best model performance for each test track on the metrics included in the main paper.

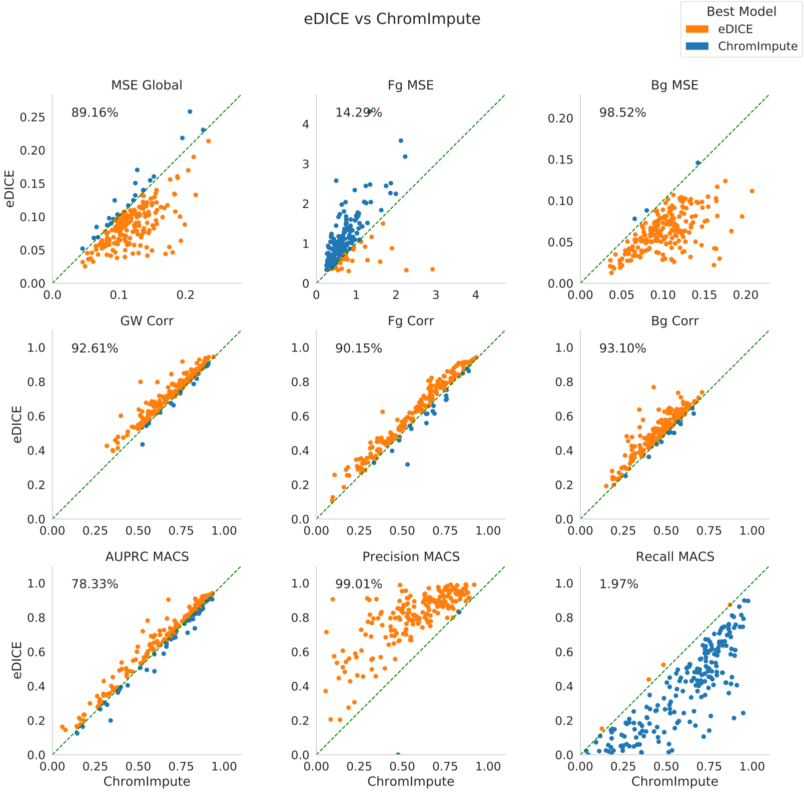


**Fig. S4** Scatterplots for the comparison of the performance of eDICE against ChromImpute for each individual test track on a subset of the evaluation metrics metric. Each point corresponds to an individual test track, whose coordinates are given by the metric of the plot for the baseline and eDICE. The percentage in the top left corner of each plot indicates the percentage of tracks on which eDICE outperform the baseline.

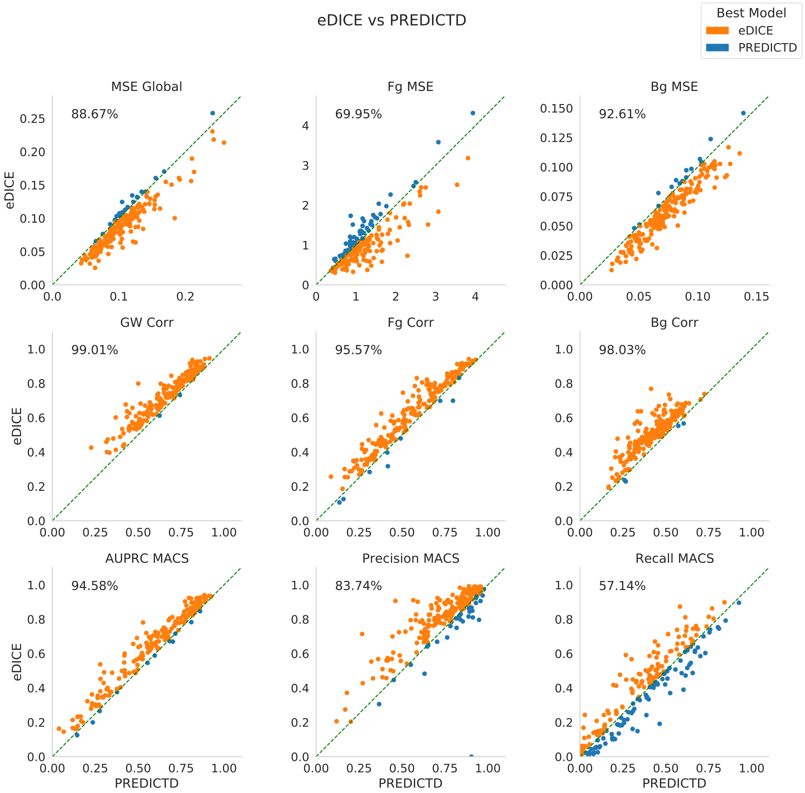


**Fig. S5** Scatterplots for the comparison of the performance of eDICE against PREDICTD for each individual test track on a subset of the evaluation metrics metric.

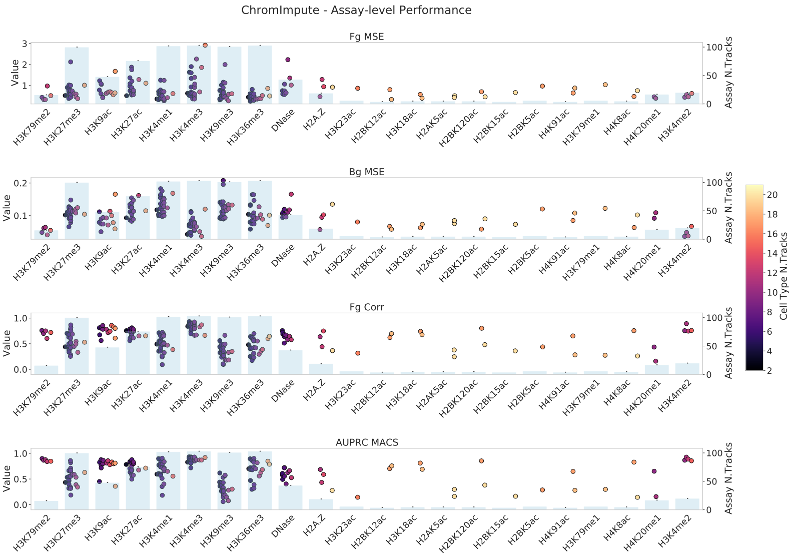


**Fig. S6** Track level performance of the imputations divided by assay for ChromImpute on the 203 test tracks from the Roadmap dataset.

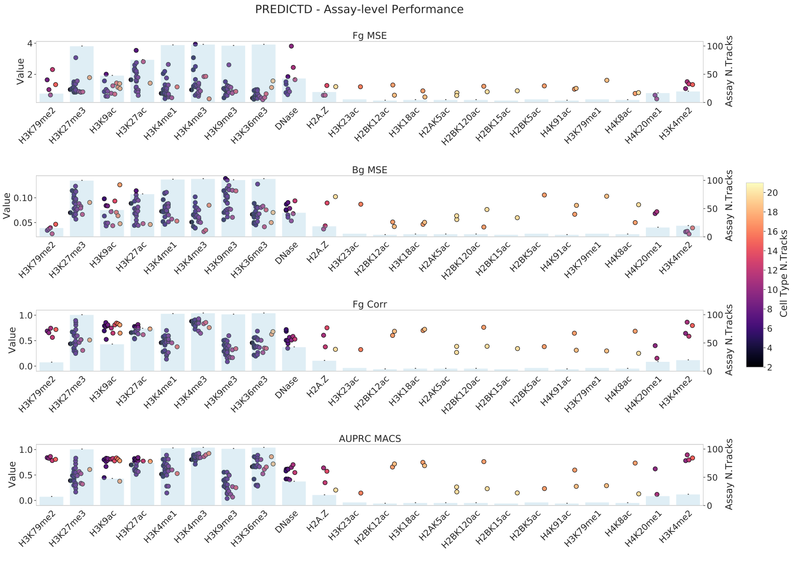


**Fig. S7** Track level performance of the imputations divided by assay for PREDICTD on the 203 test tracks from the Roadmap dataset.

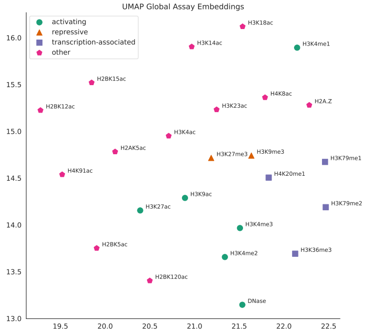


**Fig. S8** UMAP projection of assay global embeddings

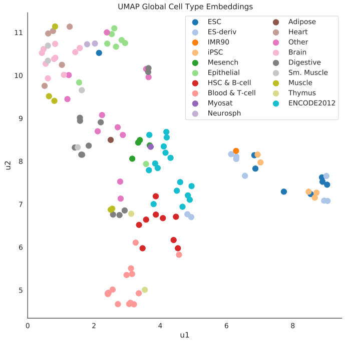


**Fig. S9** UMAP projection of cell type global embeddings

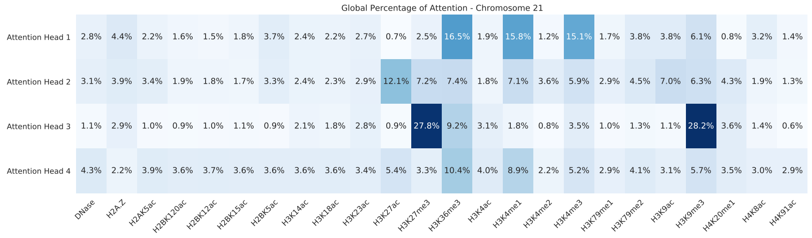


**Fig. S10** Percentage of attention that the 4 attention heads dedicate to each assay over the entire chromosome 21.

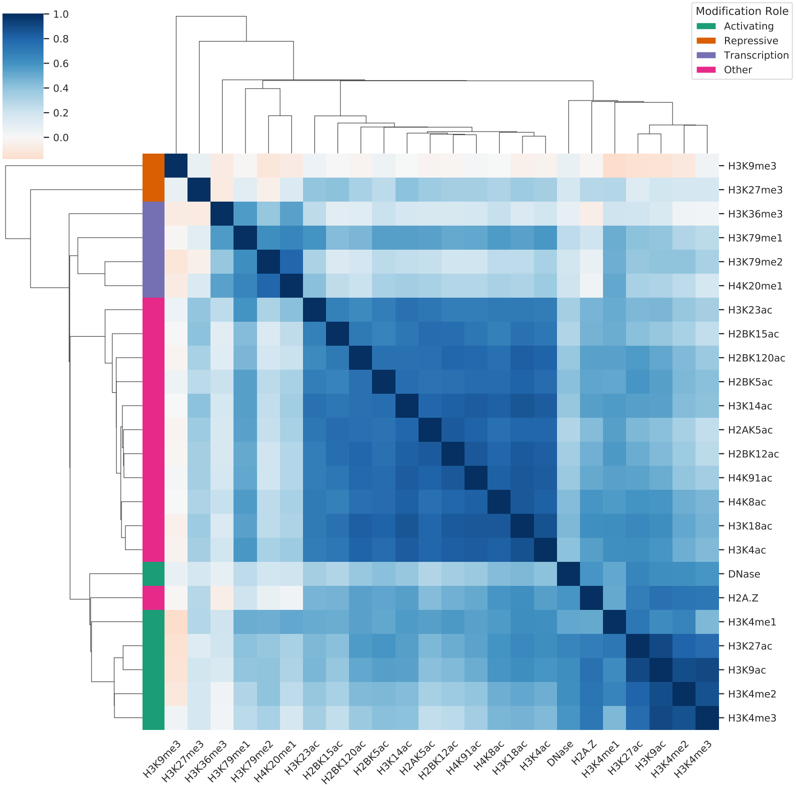


**Fig. S11** Hierarchical clustering of the correlation matrix between epigenetic marks, calculated using the average tracks over all cell types.

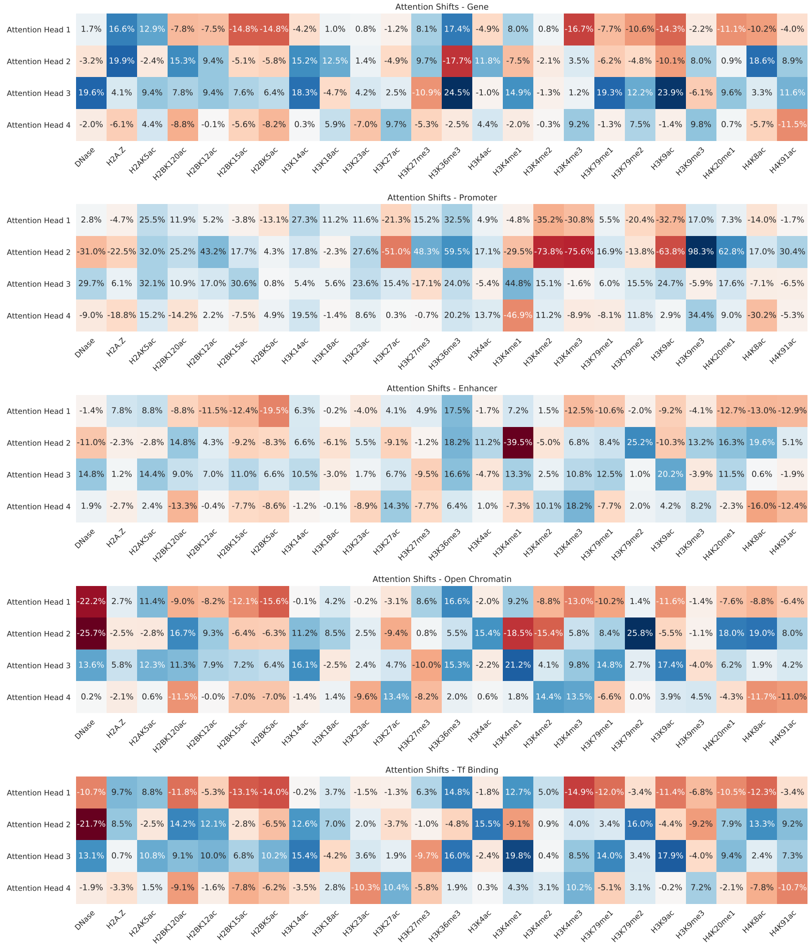


**Fig. S12** Differential percentage of attention that the 4 attention heads dedicate to each assay over the annotated portions of chromosome 21.

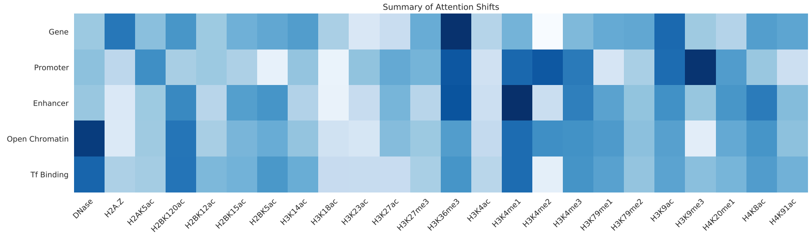


**Fig. S13** Visual summary of the attention shifts for each assay in annotated regions of the genome.

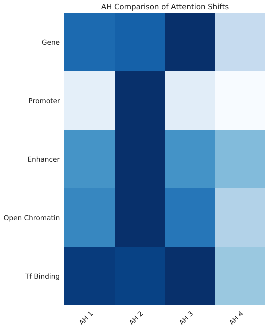


**Fig. S14** Visual summary of the attention shifts for each attention head in annotated regions of the genome.

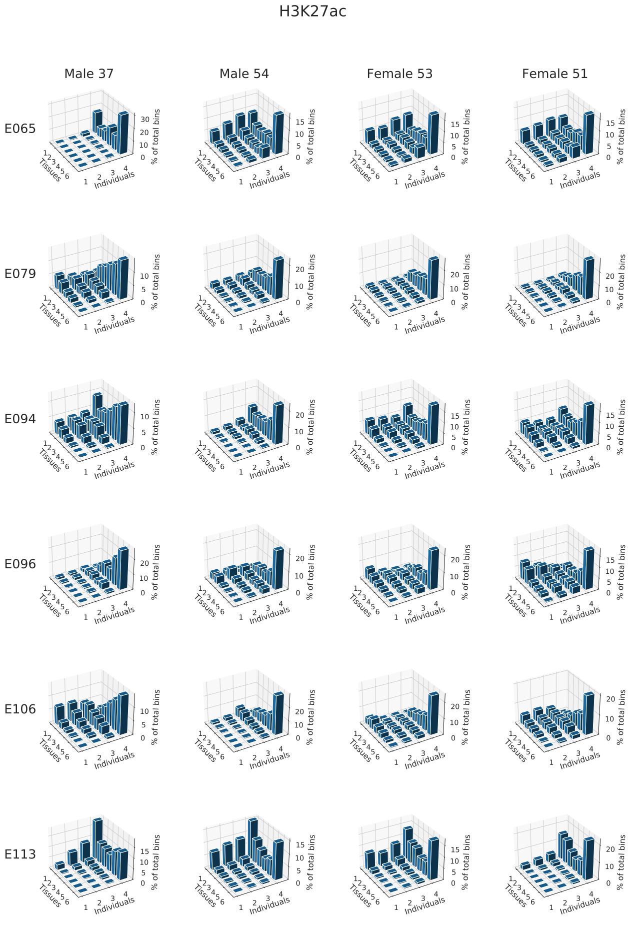


**Fig. S15** 3D Histograms representing between how many tissues and individuals each enriched bin is shared for H3K27ac tracks.

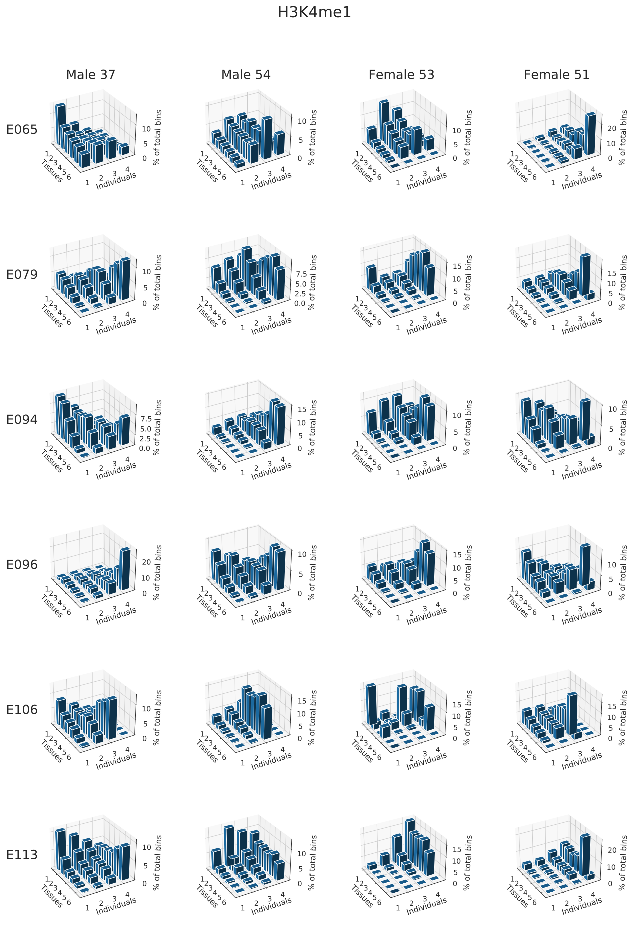


**Fig. S16** 3D Histograms representing between how many tissues and individuals each enriched bin is shared for H3K4me1 tracks.

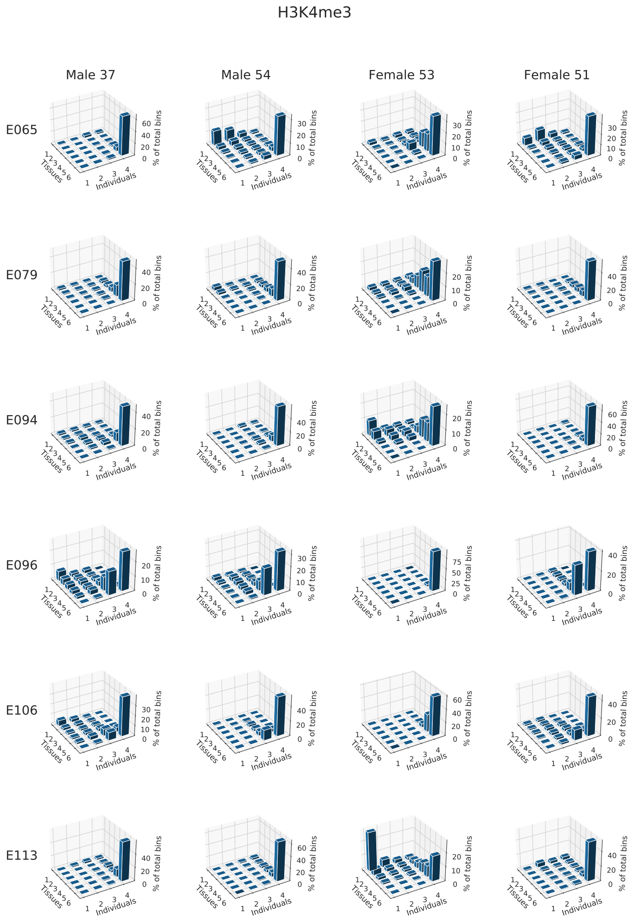


**Fig. S17** 3D Histograms representing between how many tissues and individuals each enriched bin is shared for H3K4me3 tracks.

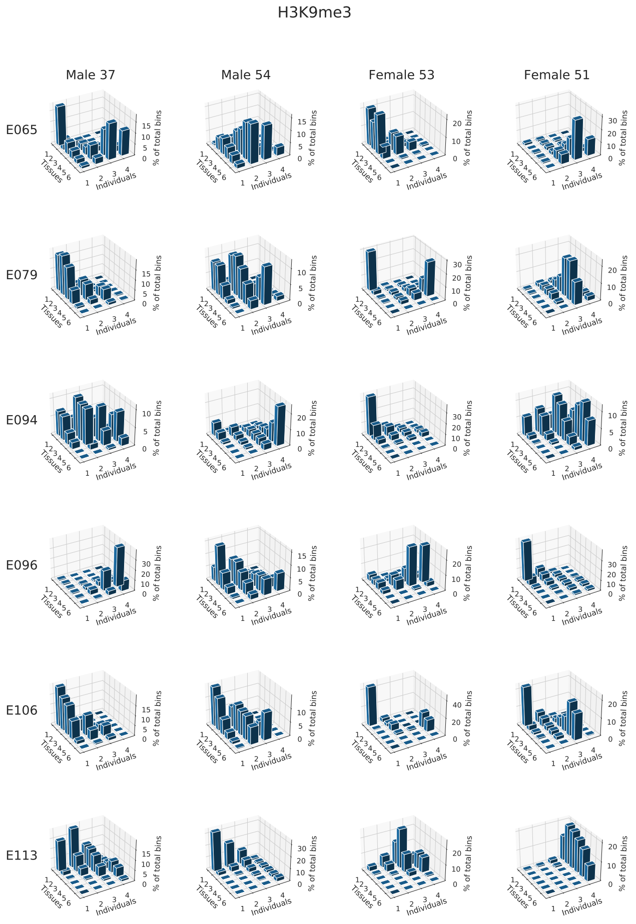


**Fig. S18** 3D Histograms representing between how many tissues and individuals each enriched bin is shared for H3K9me3 tracks.

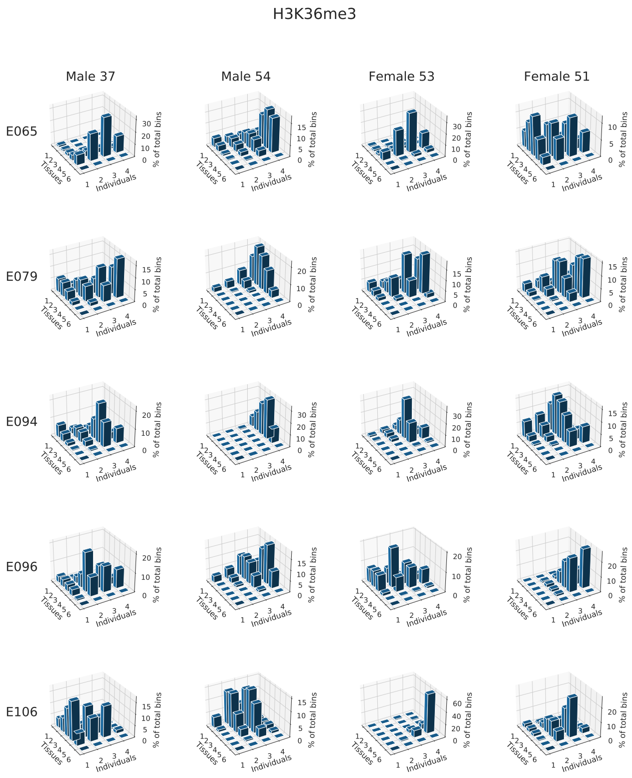


**Fig. S19** 3D Histograms representing between how many tissues and individuals each enriched bin is shared for H3K36me3 tracks.

### S2 Supplementary Tables

**Genome-Wide Signal Reconstruction**

| | GW Corr | MSE Global |
|---|---|---|
| AVG | 0.563 ± 0.02 | 0.159 ± 0.008 |
| ChromImpute | 0.697 ± 0.019 | 0.119 ± 0.005 |
| PREDICTD | 0.669 ± 0.02 | 0.106 ± 0.005 |
| eDICE | **0.735 ± 0.018** | **0.091 ± 0.005** |

**Background Bins Reconstruction**

| | Bg Corr | Bg MSE |
|---|---|---|
| AVG | 0.311 ± 0.015 | 0.124 ± 0.005 |
| ChromImpute | 0.455 ± 0.016 | 0.102 ± 0.004 |
| PREDICTD | 0.418 ± 0.016 | 0.074 ± 0.003 |
| eDICE | **0.503 ± 0.016** | **0.063 ± 0.003** |

**Foreground Bins Reconstruction**

| | Fg Corr | Fg MSE |
|---|---|---|
| AVG | 0.467 ± 0.028 | 1.328 ± 0.097 |
| ChromImpute | 0.571 ± 0.028 | **0.73 ± 0.057** |
| PREDICTD | 0.548 ± 0.029 | 1.189 ± 0.087 |
| eDICE | **0.614 ± 0.028** | 1.041 ± 0.082 |

**Obs. MACS vs Imp. Bins Classification**

| | AUPRC MACS | AUROC MACS |
|---|---|---|
| AVG | 0.45 ± 0.029 | 0.931 ± 0.006 |
| ChromImpute | 0.625 ± 0.032 | 0.967 ± 0.004 |
| PREDICTD | 0.592 ± 0.032 | 0.96 ± 0.005 |
| eDICE | **0.651 ± 0.031** | **0.969 ± 0.004** |

**Obs. MACS vs Imp. MACS Classification**

| | F1 MACS | MCC MACS | Precision MACS | Recall MACS |
|---|---|---|---|---|
| AVG | 0.41 ± 0.024 | 0.415 ± 0.023 | 0.451 ± 0.027 | 0.451 ± 0.03 |
| ChromImpute | **0.544 ± 0.027** | **0.554 ± 0.025** | 0.574 ± 0.029 | **0.603 ± 0.033** |
| PREDICTD | 0.453 ± 0.032 | 0.485 ± 0.029 | 0.739 ± 0.025 | 0.371 ± 0.031 |
| eDICE | 0.481 ± 0.034 | 0.52 ± 0.029 | **0.803 ± 0.023** | 0.392 ± 0.033 |

**Table S1** Performance metrics for the imputation of the 203 test tracks on chromosome 21. In bold we highlighted the best performance for each metric, as well as the models that failed to reject the null hypothesis of a shared mean in a two-sided paired t-test with a signficance level of 0.01.

| Tissue | Assay | ENCDO845WKR (male 37 years old) | ENCDO451RUA (male 54 years old) | ENCDO793LXB (female 53 years old) | ENCDO271OUW (female 51 years old) |
|---|---|---|---|---|---|
| ascending aorta | H3K27ac | | | ENCFF472VCY | ENCFF002VCM |
| | H3K36me3 | | | ENCFF529EEQ | ENCFF602IDJ |
| | H3K4me1 | | | ENCFF971TZV | ENCFF843GKY |
| | H3K4me3 | | | ENCFF229NZH | ENCFF383JQQ |
| | H3K9me3 | | | ENCFF616UAG | ENCFF644UVB |
| thoracic aorta | H3K27ac | ENCFF652JBU | ENCFF344YDV | | |
| | H3K36me3 | ENCFF700ZLW | ENCFF556REM | | |
| | H3K4me1 | ENCFF141KEC | ENCFF388BWY | | s |
| | H3K4me3 | ENCFF327VEH | ENCFF723SJI | | |
| | H3K9me3 | ENCFF221YER | ENCFF198OYC | | |
| esophagus muscularis mucosa | H3K27ac | ENCFF969PNR | ENCFF437XOA | ENCFF282WYM | ENCFF707HDC |
| | H3K36me3 | ENCFF476ILL | ENCFF567TNH | ENCFF723KNC | ENCFF861YVB |
| | H3K4me1 | ENCFF295FVH | ENCFF478GEN | ENCFF067DEH | ENCFF877IZN |
| | H3K4me3 | ENCFF280VIW | ENCFF979ERR | ENCFF156BAJ | ENCFF636QVZ |
| | H3K9me3 | ENCFF501FLV | ENCFF380LRK | ENCFF885KSL | ENCFF729RCC |
| stomach | H3K27ac | ENCFF440VXQ | ENCFF241NFR | ENCFF134NHD | ENCFF140JKH |
| | H3K36me3 | ENCFF302OSD | ENCFF414FQR | ENCFF428LSW | ENCFF441GAP |
| | H3K4me1 | ENCFF140RPG | ENCFF034VHX | ENCFF411ZSD | ENCFF131UFO |
| | H3K4me3 | ENCFF175WCH | ENCFF221JKN | ENCFF309DCR | ENCFF468UJW |
| | H3K9me3 | ENCFF285GPE | ENCFF765FRJ | ENCFF674DMF | ENCFF554RBC |
| upper lobe of left lung | H3K27ac | ENCFF204PWD | ENCFF305XCG | ENCFF071QUR | ENCFF178DCM |
| | H3K36me3 | ENCFF549FDA | ENCFF572VAG | ENCFF267DPL | ENCFF700BQU |
| | H3K4me1 | ENCFF076BNQ | ENCFF614CUN | ENCFF777AOP | ENCFF344GZI |
| | H3K4me3 | ENCFF898DYH | ENCFF526RLU | ENCFF887IUV | ENCFF578OBF |
| | H3K9me3 | ENCFF672QOH | ENCFF583KBZ | ENCFF076RRH | ENCFF317KCA |
| sigmoid colon | H3K27ac | ENCFF669TSI | ENCFF507VUO | ENCFF156QWL | ENCFF497GAF |
| | H3K36me3 | ENCFF284VTI | ENCFF312AHC | ENCFF577WWK | ENCFF744WIO |
| | H3K4me1 | ENCFF686SIG | ENCFF427WMO | ENCFF126IMK | ENCFF113YOI |
| | H3K4me3 | ENCFF892GNP | ENCFF445QHF | ENCFF905JEN | ENCFF135XHH |
| | H3K9me3 | ENCFF811EKT | ENCFF925APW | ENCFF782WTP | ENCFF477NHW |
| spleen | H3K27ac | ENCFF230ZQJ | ENCFF786NLG | ENCFF735GCH | ENCFF702AJP |
| | H3K4me1 | ENCFF873LAX | ENCFF049HBP | ENCFF402YZU | ENCFF599IBO |
| | H3K4me3 | ENCFF422ZXF | ENCFF189KXE | ENCFF656AAS | ENCFF462VQE |
| | H3K9me3 | ENCFF952NOQ | ENCFF418KYC | ENCFF447CEL | ENCFF088KFT |

**Table S2** Accession codes for the 116 selcted tracks from the ENTEx dataset.

| ENTEx Tissue | Roadmap EID |
|---|---|
| ascending aorta | E065 |
| thoracic aorta | E065 |
| esophagus muscularis mucosa | E079 |
| stomach | E094 |
| upper lobe of left lung | E096 |
| sigmoid colon | E106 |
| spleen | E113 |

**Table S3** Mapping of tissues between the ENTEx dataset and the Roadmap cell type IDs.

### S3 Results

#### S3.1 Evaluation Strategy

Measuring the performance of an imputation model is far from trivial. Properly designing an evaluation strategy is crucial to ensure that the results reflect the workings of the models and that the comparison to a selected set of baselines aligns with the use case intended for the model.

Ernst and Kellis [S1] validated the performance of ChromImpute with a leave-one-out (LOO) strategy, where all tracks short of the one specific target test track are used during different training instances. This strategy provides a substantial prediction advantage due to the optimal number of training data sets. However, it may not give the best assessment of the method's generalisation ability [S2]. In contrast, the authors of PREDICTD and Avocado have analysed the imputed values for a hold-out set of test tracks using 5-fold cross-validation. This procedure has also been adopted in a recent ENCODE Imputation Challenge, and we will do the same in this work. The split of the data is identical to one of the folds used in [S3]. Supplementary Figure S1 depicts a data matrix showing which tracks from the Roadmap consortium were part of the training, validation and test sets. Quality control of the Roadmap Consortium guarantees a minimum standard for all data sets [S4].

In previous work, including the Encode Imputation Challenge, training, validation and test tracks are processed following established computational pipelines[1]. This pipeline includes a pooling step for biological and technical replicates and subsampling to maximum library size. Notably, the MACS2 peak caller is applied to calculate the statistical significance of enrichment at each base pair in the genome. The resulting genome-wide signal tracks containing the statistical significance of enrichment (i.e., the $-log10$ p-values) at each base pair in the genome have been used both as input training data and to validate imputations [S3]. Recently, however, it has been demonstrated that MACS2 outputs biased P-values and false discovery rate estimates that can be many orders of magnitude too optimistic [S5]. This systematic bias raises several problems; for instance, there is no sound basis for choosing a p-value cut off for reporting results. Additionally, as the introduced bias depends on the ChIP-Seq data from which the p values are computed, there can be "distributional shifts" between training and test sets, as observed in the Encode Imputation Challenge.

To reduce the impact of possible systematic biases, we employ a varied set of performance metrics, each aiming to capture different facets of the imputation task. For example, comparing models using Area-Under-Curve metrics ensures that choosing a specific p-value cutoff is not the determining factor in ranking the imputation models.

#### S3.2 Performance Metrics

An appropriate choice of metrics is essential for robust comparisons between imputation models. The most immediate and intuitive approach from a machine learning point of view is to measure the ability of a model to reconstruct genome-wide signals (i.e. corrected -log10 p' values). Commonly used measures include genome-wide mean-squared-error (GW MSE) and genome-wide Pearson correlation (GW Corr) [S3]. Importantly, however, most Histone modifications are only significantly enriched on a small subset of genomic bins. The remaining bins have low background signals, which are strongly determined by experimental conditions including unspecific binding of the antibody. As these background regions are over-represented on the genome they easily dominate genome-wide assessment measures.

Durham et al. [S6] therefore defined additional performance measures including MSE1obs, which measures the MSE on the top 1% of the positions according to ChIP-seq signal; and MSE1imp measures the MSE on the top

[1] e.g. https://github.com/ENCODE-DCC/chip-seq-pipeline2.

1% of the positions according to the imputed signal. On the other hand, outside the imputation literature, it is a common strategy to analyse histone modification measurements by focusing on foreground regions which are identified using Peak callers such as MACS2 [S7]. In this case, significant enriched regions are merged if they are close together and fluctuations below a given threshold are tolerated within a foreground peak if they are small enough. Additionally, regions of significant enrichment have to have a minimum length (at least as big as the fragment length) in order to be called foreground peaks. Peak callers therefore provide a more robust segmentation of the genome into foreground and background regions. Here we follow a hierarchical analysis to evaluate the imputations: First, we test the ability of the model to correctly classify genomic regions into foreground and background regions (e.g. [S8]). The signal reconstruction potential of the models is then separately analysed for bins categorised as foreground and background. For the foreground regions, we are additionally interested in how well the peak intensities are predicted (e.g. [S9]), and whether the shape of the signal in the peak is well captured by the prediction. (e.g.[S10, S11]).

#### S3.3 Validation on the Roadmap Reference Epigenome

Table S1 presents the numerical results for the imputation of the test tracks of the Roadmap dataset. For each metric we highlighted the best-performing model, as well as the competitors for which a 2-sided t-test failed to reject the null hypothesis with a significance threshold of 0.01.

Comparing eDICE to the baselines at the assay level (Figure S2) and at the individual track level (Figures S3, S4, and S5) highlights that the reported improvements are consistent across the board, with the notable exception of four metrics in which ChromImpute ourperforms eDICE (Fg MSE, Precision MACS, F1 MACS, and MCC MACS). For all four of them, the underlying issue is that factorisation models such as eDICE and PREDICT consistently underestimate the height of peaks, leading to reduced detection of enriched regions. As a consequence, eDICE is more conservative than ChromImpute in its prediction, a fact that is mirrored in the Recall MACS metrics.

Different assays present different overall imputation performance. For example, in the evaluation of eDICE, H3K9me3 and H3K27me3 displayed consistently suboptimal performance. Figures S6 and S7 highlight how this tendency is not exclusive to eDICE but shared by the baseline models.

#### S3.4 Differential peak analysis

Differential peak analysis aims to detect meaningful differences between biological samples and typically involves multiple replicates in each group of interest. The inclusion of multiple samples enables the discrimination of natural variability within cell types from differentially enriched regions caused by the biological differences between tissues.

To analyse the use of imputed signals for detecting differential enrichment, we chose the H3K9ac assays for cell types E025 (Adipose Derived Mesenchymal Stem Cell Cultured Cells) and E052 (Muscle Satellite Cultured Cells) as samples for a case study. These two epigenomic tracks are part of the hold-out test set, share a comparable number of tracks for the two tissues in the training, validation, and test sets, and have 4 and 3 measured replicates, respectively.

We tested the use of imputations for differential analysis with the DiffBind package [S12]. DiffBind requires specifying the peak set and providing the aligned reads for treatment and control in a sample sheet. For the measurement group, we use the data from the Roadmap consortium, which includes the aligned reads for each replicate in *.tagAlign* format and the peaks called with MACS2 in *.narrowPeak* format. In addition, each replicate is associated with a cell-type-specific control track. The aligned *.tagAlign* files are converted to BAM format with BedTools [S13].

For the imputed tracks, we devised a procedure to simulate replicates from the predicted p-value signal. Such a procedure is an approximation that requires several assumptions, yet it suffices to provide a proof of concept for using imputation models in bioinformatics software pipelines. Nevertheless, for such applications, future imputation models would likely need to focus on modelling not only the average value of a signal but also model the uncertainty involved.

To simulate replicates using p-value signals, we assume that the read counts at each genomic position follow a negative binomial distribution, a common model used for sequencing data [S14–S16]. To draw samples from this distribution, we need to estimate two parameters, mean and dispersion.

We estimate the mean parameter by inverting the procedure used to generate the p-value tracks. The p-value signal results from the Survival Function of a background Poisson model. The genomic-position specific mean parameter of the background distribution is calculated from tissue-specific control samples available in the Roadmap dataset. Using the parameters of the background distribution, the read counts for a given track are obtained through the Inverse Survival Function (ISF).

To infer the dispersion parameters for each cell type we made use of the software package DESeq2 [S15]. Specifically, DESeq2 is capable of fitting a function that, given a the mean value as input, can output the dispersion value. We selected tracks from the training set for H3K9ac whose tissues present sufficient similarity to the target tissues (E023 for E025 and E107-E108 for E052). These selected tracks are used to estimate the mean-dispersion functions, which are then used on the previously estimated mean parameters to generate bin-wise dispersion parameters.

Given these bin-specific parameters derived from the imputed signals, we generate three samples per cell type by sampling from the negative binomial distribution.

The MACS[S7] command *bdgpeakcall* was used to generate the peak set on the p-value track for each cell type in both measurement and simulated replicates. Next, the consensus peakset for all samples was estimated using DiffBind. ENCODE blacklisted regions[S17] were removed from the consensus peakset. Aggregated counts for these consensus peaks were computed for both measurement and simulated replicates, and were directly read into the DiffBind DBA object with *dba.peakset* function. Using a False Discovery Rate (FDR) threshold of 0.05, we determined the differentially enriched peaks for both measurements and imputations.

##### S3.5 Individual Epigenomes

For the case study of personalised imputations, we downloaded a set 29 tissue-assay combinations shared by all four individuals in the ENTEx dataset, for a total of 116 tracks. The accession codes for these tracks are listed in Table S2.

The tissues and assays represented in this set of tracks were chosen to overlap as much as possible with those present in the Roadmap dataset, in order to enable the use of a transfer learning framework. This resulted in the selection of 5 histone modifications that are shared with the Roadmap set, and 6 tissues, that present an approximate overlap (Table S3). Two tissues (ascending aorta and thoracic aorta) were combined into a single Roadmap ID. The imprecise mapping of cell types is an added source of systematic noise that imposes considerable challenges for the imputation of individual epigenomes.

Each p-value track was binned at a 25 base pairs resolution, following the processing steps used for the Roadmap dataset.

#### S4 Interpreting the model

##### S4.1 Global Embeddings

eDICE learns global embeddings that capture the high-level relationships between assays and cell types, which we can display through a UMAP [S18] projection.

Epigenetic modifications that are generally associated with the same functionality tend to cluster together in the learned representations (Figure S8). Assigning global roles to epigenetic modifications neglects much of the nuance of the regulatory mechanisms of the genome. Nonetheless, we can consider some general patterns.

H3K9ac, H3K4me2, H3K4me3 are modifications typically associated with active promoters [S19, S20], while H3K4me1 and H3K27ac (and partially H3K4me2) are linked with active enhancers [S20, S21]. We classify these marks, together with the DNAse-seq assay [S22], under the broad label of *activating* modifications.

H3K27me3 is an antagonist to H3K27ac and is broadly associated to gene repression together with H3K9me3 [S23, S24].

H3K36me3, H4K20me1, H3K79me1 are linked to exons [S25], and we consider these modifications under the label of *transcription-associated* together with H3K79me2, which is associated with active gene bodies and alternative splicing [S26, S27].

The partitioning of the epigenetic marks reflects the correlation patterns that emerge from the observed data. We calculated the average tracks for each assay across all cell types and performed agglomerative hierarchical clustering on these tracks. The results, summarised in Supplementary Figure S11, show that modifications with shared global functions tend to cluster together.

Likewise, eDICE is capable of learning the broad similarities between tissue types through the global cell embeddings represented in Figure S9.

#### S4.2 Measures of Attention

Inspired by the analysis of attention mechanisms in protein language models [S28], we define the portion of attention that each attention head $h$ dedicates to a specific assay $a$ in the set of observed assays $A$ over a portion $g$ of the genome as:

$$p_h(\alpha) = \sum_{g \in G} \sum_{a \in A} w_{a\alpha}^{(g,h)} / \sum_{g \in G} \sum_{a \in A} \sum_{a' \in A} w_{aa'}^{(g,h)} \tag{S1}$$

where $w_{a\alpha}^{(g,h)}$ denotes the attention weight for the key assay $\alpha$ given the query assay $a$ in the genomic bin $g$ for attention head $h$. Within the formalism of key-value-query attention in Transformer models, this definition of portion of attention corresponds to the average attention given to each key. Due to the softmax function used in the calculation of the attention weights in the scaled dot-product attention, the portion of attention adds up to one over the space of all assays for a single attention head, which makes it suitable to be expressed as a percentage of the total attention of the attention head.

To evaluate how the state of the attention blocks changes in correspondence of functional regions of the genome, we calculated of the attention percentage while restricting the genomic bins to the aforementioned annotated regions and evaluated the relative difference in the percentage of attention dedicated to the assays compared to the global pattern in the chromosome. We define this *attention shift* for a given region $R$ of the genome as:

$$d_h^{(R)}(\alpha) = \frac{p_h(\alpha)|_{G \equiv R} - p_h(\alpha)}{p_h(\alpha)} \tag{S2}$$

where R is any type of annotated region of the genome, e.g. promoters or enhancers.

#### S4.3 Interpreting the Attention Weights

Attention models assign a weight to the relationship between elements in an input set or sequence, which is often considered a proxy to interpret the underlying interactions between said elements. This area of research is mainly active within the application of Transformer models to Natural Language Processing (NLP) tasks [S29, S30], with a strong focus on the visualisation of the attention weights (e.g. [S31]).

Despite this excitement, recent works have begun to highlight the limitations of attention as an explanation method, showing that in specific contexts, attention weights are at best noisy predictors of the importance of input components for a model [S32] or outright misleading and weak to adversarial attacks [S33]. Although these studies present some critical limitations, such as the focus on NLP models or the comparison to alternative measures of feature importance which are not necessarily valid as "ground truth", we believe caution in the analysis of attention weights to be warranted. Adding these considerations to the non-deterministic result of training attention models, we consider the analysis presented here as a diagnostic exploration rather than a detailed claim on the biology examined by the model.

The results presented are derived from the same instance of eDICE used for the reference epigenome validation. Although the details vary due to the stochastic training of the attention block, we observed that the conclusions drawn tend to hold across training instances.

We specifically focus on the attention block used for the contextualisation of the assay embeddings. The attention measures employed (detailed in Supplementary Section S4.2) are based on averages of attention weights, which paint a higher-level picture than comparisons of individual coefficients.

To obtain a general overview of the eDICE model, we calculated the percentage of attention given by each head to each assay for the entire chromosome 21. These results, displayed in Figure S10, highlight how at least two of the attention heads consistently dedicate a significant portion of attention to just a few histone modifications. These marks are among the core modifications extensively mapped across many cell types in the Roadmap dataset, i.e. H3K27ac, H3K36me3 H3K9me3, H3K4me1, and H3K4me3.

Interestingly, this phenomenon proved quite consistent over different experiments, with minor variations such as focusing on H3K27me3 rather than H3K27ac. Overall, this result suggests that the model relies extensively on marks for which a large amount of information is present during training.

To further explore the workings of the attention layer in eDICE, we gathered genomic annotations for chromosome 21 from the Ensembl [S34] version 104 GRCh37 assembly for protein-coding gene bodies, enhancers, promoters, open chromatin, and transcription factors binding sites. We calculated how the percentage of attention shifts within these functional regions of the genome and displayed the results in Figure S12.

Overall, these attention shifts reveal known patterns from the literature yet present unexplained peculiarities, such as the significant shift within enhancer regions for H3K4me1, a known chromatin hallmark for these functional regions. Intuitively, one would expect the attention weight for H3K4me1 to grow within enhancers, given its biological importance, yet we observe the opposite, especially for attention head 2. Nevertheless, this pattern is not consistent for such a setting: H3K36me3, a histone modification enriched on the gene body region associated with active gene transcription [S35], shows a positive average attention shift within protein-coding genes.

Much remains to be understood about the workings of attention models. Therefore, we limit our analysis to the qualitative consideration that the larger attention shifts within functional regions correspond to known epigenetic marks that characterise said regions. Examples of these patterns include the DNase shifts for open chromatin regions [S36], the shifts in DNase and H3K4me1 for transcription factor binding sites [S37], and the shifts within promoters for H3K4 methylation [S38], H3K9ac, H3K27ac, and their antagonists H3K9me3 and H3K27me3 [S19].

We include two high-level visual summaries of the attention shifts in Figures S13 and S14, in which the absolute value of the attention shifts was summed up over the attention heads and the assays, respectively, and then 0-1 normalised for each annotated category. The resulting heatmaps provide at a glance an overview of which assays present the most considerable relative changes and which attention heads are significantly affected within functional regions of the genome.

Interestingly, it appears that the attention heads specialise at least partially, with attention head 2 being the most affected in promoters, enhancers, and open chromatin regions. In contrast, attention head 3 presents the most significant shifts for protein-coding gene bodies and transcription factor binding sites. The scarce shifts for attention head 4 also indicate that this head is not contributing significantly to the final prediction, a known phenomenon in the use of multi-headed attention models, and could be a candidate for pruning [S39].

### S5 ENCODE imputation challenge model

An early version of our model was used as the basis of our third-placed entry in the 2019 ENCODE Imputation Challenge. This model follows Avocado in adopting the factorised structure of Equation 1, but instead of directly parametrising genomic bin representations $\mathbf{b}_k$, computes non-linear, low dimensional transformations of the input signal at each bin,

$$\hat{Y}_{ijk} = g_\theta(\mathbf{c}_i, \mathbf{a}_j, f_\phi(\text{vec}(Y^k))) \quad , \tag{S3}$$

where $\text{vec}(Y^k)$ is a fixed-size vector whose entries are the signal values for all observed tracks at the $kth$ bin together with zeros for all unobserved tracks. In practice the functions $g_\theta$ and $f_\phi$ are both implemented with two-layer MLPs, and the $\mathbf{c}_i$ and $\mathbf{a}_i$ are learned cell and assay embeddings, equivalent to the 'global embeddings' $\mathbf{u}_{c_i}$ and $\mathbf{u}_{a_i}$ above (Equations 4 and 6). This model can be seen as a kind of denoising autoencoder in which the decoder has a factorised structure, inherited from Avocado, allowing it to impute signal values at previously unobserved cell-assay combinations. In contrast our full model is fully factorised, replacing a single global transformation of the input signal with cell- and assay-dependent transformations of the input signal, learned via self-attention (Equation 2).

The challenge model is used as a strong baseline in our experiments, since it has demonstrated competitive performance with state-of-the-art methods including Avocado in the blind-test setting of the ENCODE Imputation Challenge.

[S34] Howe, K.L., Achuthan, P., Allen, J., Allen, J., Alvarez-Jarreta, J., Amode, M.R., Armean, I.M., Azov, A.G., Bennett, R., Bhai, J., Billis, K., Boddu, S., Charkhchi, M., Cummins, C., Da Rin Fioretto, L., Davidson, C., Dodiya, K., El Houdaigui, B., Fatima, R., Gall, A., Garcia Giron, C., Grego, T., Guijarro-Clarke, C., Haggerty, L., Hemrom, A.,

Hourlier, T., Izuogu, O.G., Juettemann, T., Kaikala, V., Kay, M., Lavidas, I., Le, T., Lemos, D., Gonzalez Martinez, J., Marugán, J.C., Maurel, T., McMahon, A.C., Mohanan, S., Moore, B., Muffato, M., Oheh, D.N., Paraschas, D., Parker, A., Parton, A., Prosovetskaia, I., Sakthivel, M.P., Salam, A., Schmitt, B.M., Schuilenburg, H., Sheppard, D., Steed, E., Szpak, M., Szuba, M., Taylor, K., Thormann, A., Threadgold, G., Walts, B., Winterbottom, A., Chakiachvili, M., Chaubal, A., De Silva, N., Flint, B., Frankish, A., Hunt, S.E., IIsley, G.R., Langridge, N., Loveland, J.E., Martin, F.J., Mudge, J.M., Morales, J., Perry, E., Ruffier, M., Tate, J., Thybert, D., Trevanion, S.J., Cunningham, F., Yates, A.D., Zerbino, D.R., Flicek, P.: Ensembl 2021. Nucleic Acids Research **49**(D1), 884–891 (2020) https://academic.oup.com/nar/article-pdf/49/D1/D884/35364073/gkaa942.pdf. https://doi.org/10.1093/nar/gkaa942

[S35] Shilatifard, A.: Chromatin modifications by methylation and ubiquitination: implications in the regulation of gene expression. Annu. Rev. Biochem. **75**, 243–269 (2006)

[S36] Boyle, A.P., Davis, S., Shulha, H.P., Meltzer, P., Margulies, E.H., Weng, Z., Furey, T.S., Crawford, G.E.: High-resolution mapping and characterization of open chromatin across the genome. Cell **132**(2), 311–322 (2008)

[S37] Robertson, A.G., Bilenky, M., Tam, A., Zhao, Y., Zeng, T., Thiessen, N., Cezard, T., Fejes, A.P., Wederell, E.D., Cullum, R., *et al.*: Genome-wide relationship between histone h3 lysine 4 mono-and tri-methylation and transcription factor binding. Genome research **18**(12), 1906–1917 (2008)

[S38] Mohn, F., Schübeler, D.: Genetics and epigenetics: stability and plasticity during cellular differentiation. Trends in Genetics **25**(3), 129–136 (2009)

[S39] Voita, E., Talbot, D., Moiseev, F., Sennrich, R., Titov, I.: Analyzing multi-head self-attention: Specialized heads do the heavy lifting, the rest can be pruned. In: Proceedings of the 57th Annual Meeting of the Association for Computational Linguistics, pp. 5797–5808 (2019)
